# A synthetic cell with integrated DNA self-replication and membrane biosynthesis

**DOI:** 10.1101/2025.01.14.632951

**Authors:** Ana María Restrepo Sierra, Federico Ramirez Gomez, Mats van Tongeren, Laura Sierra Heras, Christophe Danelon

## Abstract

The emergence, organization, and persistence of cellular life are the result of the functional integration of metabolic and genetic networks. Here, we engineer phospholipid vesicles that can operate three essential functions, namely transcription-translation of a partial genome, self-replication of this DNA program, and membrane synthesis. The synthetic genome encodes six proteins and its compartmentalized expression produces active liposomes with distinct phenotypes demonstrating successful module integration. Our results reveal that genetic factors exert a stronger control over DNA replication and membrane synthesis than metabolic crosstalk or module co-activity. By showing how genetically encoded functions derived from different species can be integrated in liposome compartments, our work opens new avenues for the construction of autonomous and evolving synthetic cells.

## Introduction

The construction of a synthetic cell from the bottom up is a grand challenge at the intersection of bioscience and engineering. Inspired by the observation of common processes in all living organisms, researchers have started to build some of the essential cellular functions, hereafter called ‘modules’, in vitro. The expanding repertoire of genetic parts and characterized biochemical networks has enabled the cell-free reconstitution of life’s fundamental mechanisms, such as the synthesis of membrane constituents (*1–3*), division related processes (*4,5*), DNA replication (*6,7*), energy regeneration (*8*), and cell-cell communication (*9*). While these studies have yielded valuable insights into the specifics of each biological module, they have not addressed the higher-ordered complexity that lies in the integration of multiple processes, in particular when the involved genetic or protein parts are derived from various organisms (*10,11*).

Three subsystems appear essential for basic cellular life: a vesicular system defining an internal machinery that synthesizes its membrane constituents, a replicable template that carries information, and a metabolic cycle that produces the molecular components (*12,13*). As a construction paradigm, we envisioned that in vitro transcription-translation (IVTT) of a synthetic DNA template using recombinant elements (PURE system) (*14*) inside phospholipid vesicles (liposomes) would constitute the scaffold onto which biological functions can be implemented to create an autonomously living synthetic cell. In contrast to other approaches which rely exclusively on purified proteins or cell lysates, our DNA-based architecture enables replication, system’s level evolution, and is constituted of well-defined components (*11*). Replication of the DNA program can be seen as the seed module priming self-maintenance and evolvability (*15*). However, an experimental demonstration of module integration directed by a synthetic self-replicating genome has remained elusive.

Here, we pinned the work for combining synthetic cell modules by constructing a novel synthetic self-replicating DNA genome, named *DNArep-PLsyn*, encoding both a DNA replication machinery (DNArep) and a phospholipid biosynthesis pathway (PLsyn). We established the conditions for in-liposome expression of *DNArep-PLsyn* with PURE system, and demonstrated the combined activities of universal cellular modules in a minimal in vitro system.

### Design and cell-free expression of a synthetic replicating genome

We constructed a synthetic DNA replication system following the design of the Φ29 genome (*16*) which consists of a linear DNA template with origin of replication sequences at each end. Previous work showed that four phage proteins – the terminal protein (TP) that functions as a replication primer, the DNA polymerase (DNAP), the single-stranded DNA binding protein (SSB), and the double-stranded DNA binding protein (DSB) – were sufficient to replicate a linear DNA in vitro (*17*). Moreover, we previously showed that expression in PURE system of a minimal Φ29-based linear replicon encoding DNAP (*p2* gene) and TP (*p3* gene) led to exponential amplification of DNA, also when the reaction was compartmentalized inside micrometer-sized liposomes (*7,15*). We here sought to integrate additional genes into this seed replication module and hypothesized that the larger synthetic genome could be replicated – and all the gene products could be synthesized – upon expression in PURE system (Fig. 1, A to D). The newly introduced genes encode four enzymes of the *E. coli* Kennedy pathway: *sn*-Glycerol-3-phosphate acyltransferase (PlsB), Lysophosphatidic acid acyltransferase (PlsC), Phosphatidate cytidylyltransferase (CdsA), and Phosphatidylserine synthase (PssA) (Fig. 1D). These enzymes catalyze the sequential conversion of oleoyl-CoA and glycerol-3-phosphate precursors into 1,2-dioleoyl-*sn*-glycero-3-phospho-L-serine (DOPS), the last intermediate for 1,2-dioleoyl-*sn*-glycero-3-phosphoethanolamine (DOPE) production. Membrane synthesis in gene-expressing vesicles can then be visualized using a PS-specific fluorescent probe (*2*). Therefore, our final linear genome, named *DNArep-PLsyn*, is flanked with Φ29 origins of replication on each end, and it encompasses six genes (two for *DNArep* and four for *PLsyn*) as individual transcription units (Fig. 1A).

**Fig. 1.**
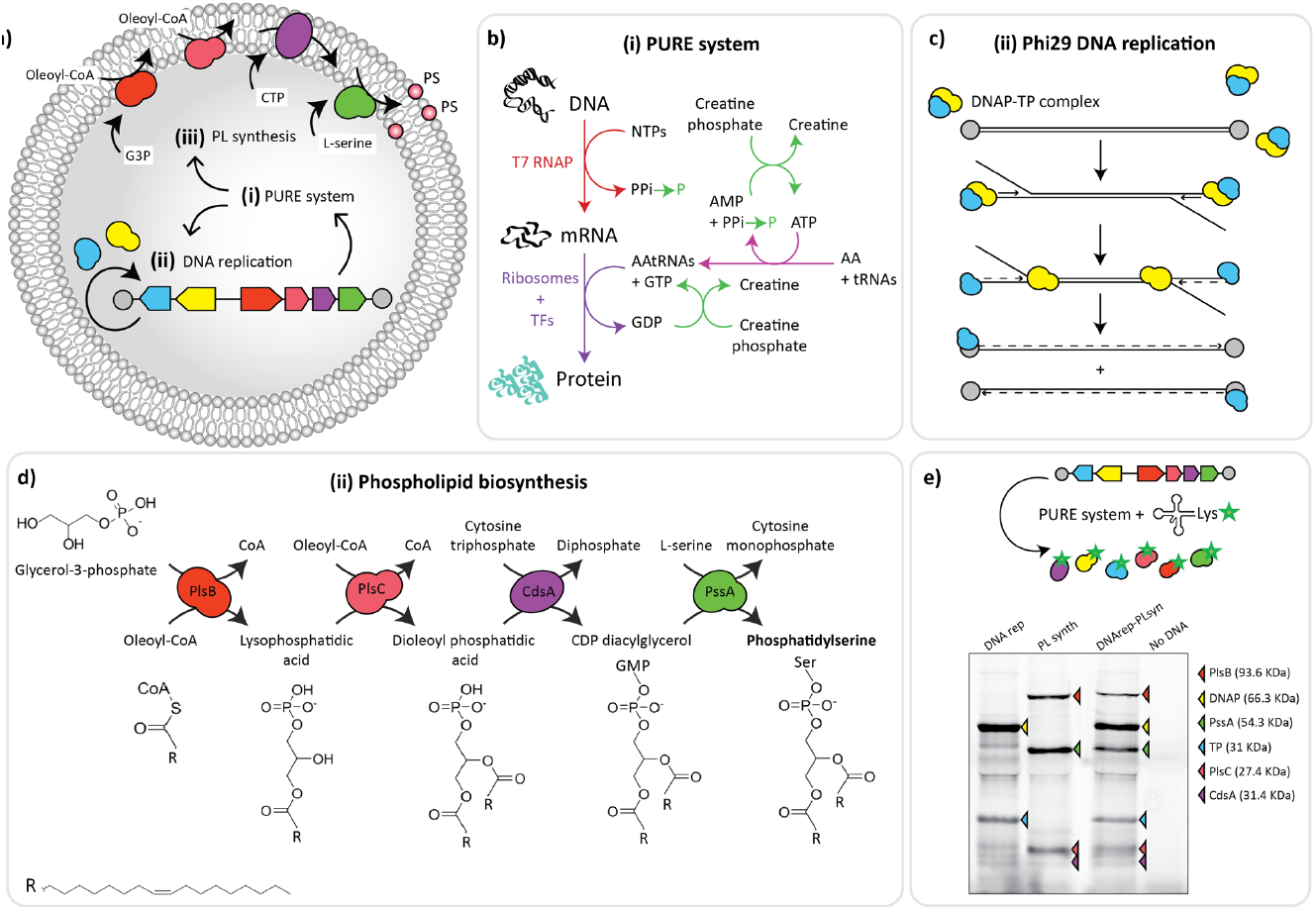
A synthetic genome encoding two cellular modules. (**A**) Synthetic vesicles with encapsulated *DNArep-PLsyn* genome and coupled transcription-translation, DNA self-replication, and phospholipid biosynthesis. (**B**) PURE system served as the main metabolic machinery for transcription and translation of DNA-encoded proteins with a creatine phosphate-based energy regeneration system. (**C**) Initiation and elongation steps of the protein-primed DNA replication mechanism from the bacteriophage Φ29. The dashed lines depict the newly synthesized strands. (**D**) A four enzyme-cascade of the *E. coli* Kennedy pathway transform*s* oleoyl-CoA and glycerol 3-phosphate into dioleoyl-phosphatidylserine (PS). (**E**) SDS-PAGE analysis of bulk IVTT reactions from the assembled *DNArep-PLsyn* template, or from the individual *DNArep* and *PLsyn* fragments. The PURE system solution was supplemented with GreenLys reagent for fluorescent labelling of the synthesized proteins (indicated with arrowheads).

To construct the *DNArep-PLsyn* synthetic genome, we iterated throughout different cloning strategies and found that template complexity (i.e., repetitive elements) often led to recombination events in *E. coli*. We then opted for an in vitro DNA assembly approach using overlapping polymerase chain reaction (PCR) to stitch the *DNArep* and *PLsyn* genetic parts (fig. S1), and a yeast-based cloning approach (fig. S2). Notably, *S. cerevisiae* yeast did not seem to pose recombination issues with repetitive regulatory sequences, unlike *E. coli*. After plasmid extraction from yeast, we generated the linear *DNArep-PLsyn* genome by PCR. In both in vitro and in-yeast DNA assemblies, we successfully obtained a linear template with the expected size (∼9,600 bp), and the sequence was validated by nanopore sequencing (fig. S1 and fig. S2).

Next, we confirmed that expression of *DNArep-PLsyn* with PURE system generates the six encoded proteins. The reaction mix was supplemented with GreenLys reagent for co-translational protein labelling. All six proteins were produced at detectable levels starting from DNA assembled in vitro or in yeast (Fig. 1E, fig. S3 and fig. S4). Interestingly, only a slight reduction of protein expression levels was observed for the *DNArep-PLsyn* template compared to the separate expression of each individual genetic module (Fig. 1E and fig. S3). This could be caused by resource sharing when the number of genes increases, but the effect was less pronounced than expected. We conclude that *DNArep-PLsyn* acts as an effective template for expressing all necessary proteins involved in both DNArep and PLsyn modules.

### Integration of DNArep and PLsyn modules inside gene-expressing vesicles

Our next aim was to evaluate and potentially optimize the simultaneous activity of the DNArep and PLsyn modules inside liposomes. Since each of the encoded modules may have a preferred reaction temperature (DNA replication works well at ∼30 °C (*7,17*), while cell-free gene expression (*18*) and phospholipid biosynthesis (*2*) are most effective at 37 °C), we decided to test different incubation temperatures. We encapsulated in liposomes the *DNArep-PLsyn* genome together with PURE system and the required substrates and cofactors for both DNArep and Plsyn, and we ran the reactions at 30°C, 34 °C, or 37 °C. After overnight incubation, we stained the DNA with the dsGreen intercalating dye (*19*) and the membrane-incorporated DOPS with the PS-specific probe LactC2-mCherry (*2*), and we analyzed the samples by flow cytometry (Fig. 2, A to D). For each fluorescent probe, we performed an intensity thresholding based on negative control samples (fig. S5), thus defining four regions of interest (ROI) in the scatter plot (Fig. 2D). Liposomes exhibiting functional DNArep (ROI 1 + ROI 2) or PLsyn (ROI 2 + ROI 4) modules were detected at all three temperatures (Fig 2, A to B), with a slightly higher occurrence for DNArep-active liposomes at 34 °C than at 30 and 37 °C (Fig. 2, B to D). Notably, a range of 0.4 to 12% of the liposomes (corresponding to ∼50 to 1,200 liposomes per sample across biological replicates at 34 °C) localized in ROI 2 indicating that both DNArep and PLsyn modules were simultaneously active (Fig. 2, C to D, and fig. S5). A larger fraction of liposomes was positive to either one of the two modules (ROI 1 or ROI 4), or was inactive (ROI 3) (Fig. 2, A, B and D). Such a heterogeneity within the same clonal (here referring to the fact that one DNA species was used) population of liposomes is also observed in single-gene expression experiments and can be attributed to uneven loading or supply of substrates or cofactors, or to varying expression levels of the genetic modules between liposomes (*20*). In addition, a significant variability across biological replicates (sample-to-sample heterogeneity) was observed. For example, the percentage of DNArep-PLsyn-positive liposomes (ROI 2) was 2.9% ± 1.3% (mean ± s.e.m) across eight biological replicates at 34 °C (fig. S5). Nonetheless, these results demonstrate that functional integration of DNArep and PLsyn modules from a synthetic genome is possible at temperatures ranging from 30 °C to 37 °C.

**Fig. 2.**
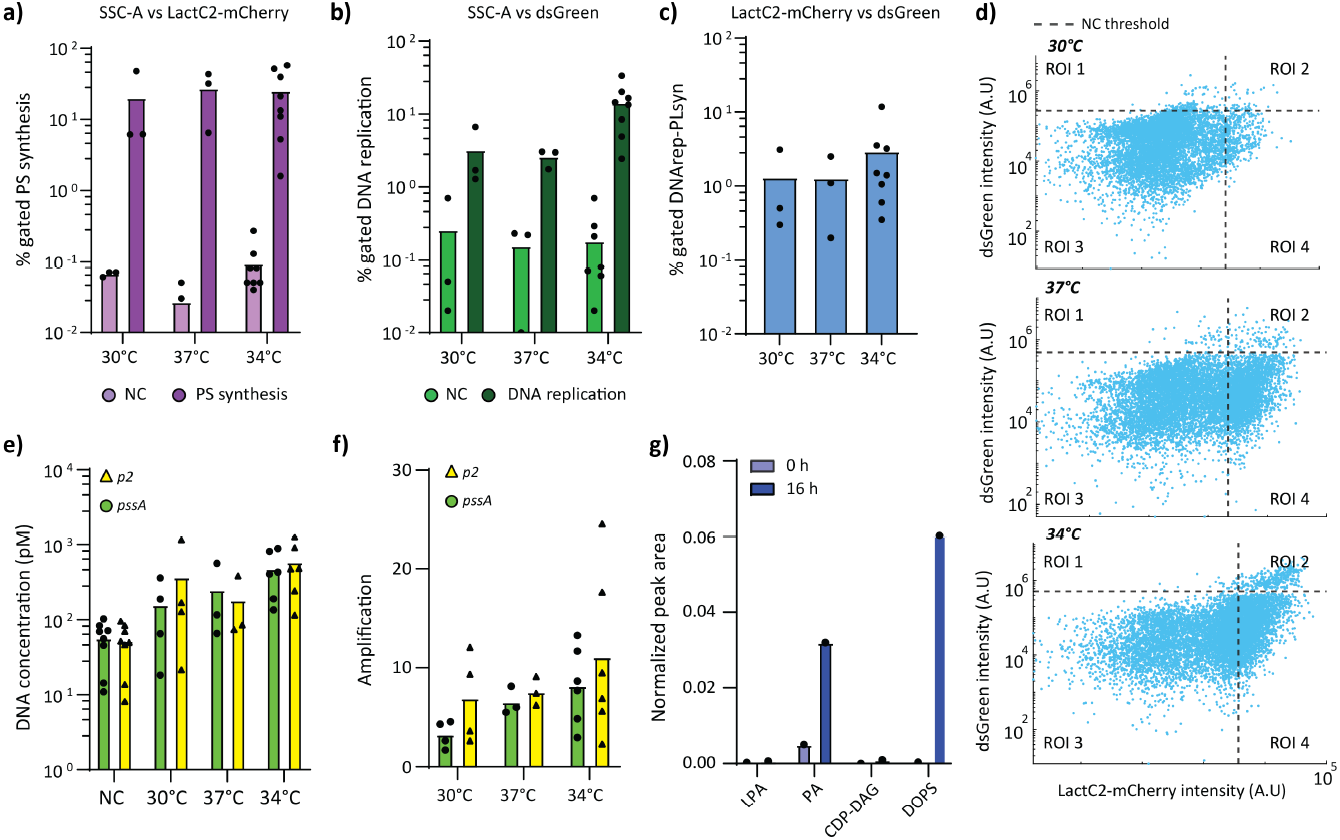
Validation of DNArep and PLsyn protein activity inside gene-expressing liposomes at different incubation temperatures. (**A**) Percentage of liposomes with active DOPS synthesis and (**B**) active DNA replication under 30, 37 and 34 °C incubation temperatures. Flow cytometry data are SSC-A vs. dsGreen for DNA replication and SSC-A vs. LactC2-mCherry for DOPS synthesis. Data points represent biological repeats and bar height the mean value. Raw data from individual replicates can be found in fig. S5. NC refers to samples, where DNA was omitted but the solutions were incubated at the indicated temperature. (**C**) Percentage of liposomes exhibiting dual dsGreen and LactC2-mCherry signals at 30, 37 and 34 °C incubation temperatures. Joint phenotype populations were selected from LactC2-mCherry vs dsGreen scatter plots. Raw data from individual replicates can be found in fig. S5. (**D**) Flow cytometry scatter plots from liposome samples displaying four regions of interest (ROI 1-4) at all tested temperatures: DNArep-active liposomes are in ROI 1, PLsyn-active liposomes in ROI 4, and liposomes with both active DNArep and PLsyn modules are in ROI 2. Vertical and horizontal dashed lines indicate intensity threshold values that have been defined using control samples (see fig. S5). Data from additional biological repeats can be found in fig. S5. (**E**) Absolute DNA quantification by qPCR of samples incubated at 30, 37, and 34 °C. qPCR target regions (∼200 bp) are from *pssA* and *p2* genes. The negative control (NC) represents calculated DNA values at initial incubation points (0 hour). (**F**) Amplification fold of *DNArep-PLsyn* DNA calculated from qPCR data in panel E: end-point (16 hours) DNA concentration / DNA concentration at time zero. Data points represent biological repeats and bar height the mean value. (**G**) LC-MS detection of DOPS and PLsyn intermediate enzymatic products before and after expression of the *DNArep-Plsyn* genome. Peak area for each compound was normalized to that of DOPG. Additional biological repeats and negative controls can be found in fig. S8.

To provide a more direct evidence of genome self-replication, we measured the concentration of DNA using quantitative PCR (qPCR). Two different sequences localized in opposite regions of the linear *DNArep-PLsyn* genome were targeted for qPCR, one in the *p2* gene and one in the *pssA* gene. The results quantitatively confirmed that all tested temperatures supported genome replication, again with a slight preference for 34 °C (Fig. 2, E to F). We further investigated whether the full-length genome was amplified (vs. shorter amplicons) by targeting all six genes by qPCR. These experiments were performed at 34 °C. Despite some variations in the concentration of replicated genes, the data showed that the entire DNA sequence between the *p3* and *pssA* genes (∼5,000 bp apart) was amplified about 10-fold (fig. S6). Small differences could arise from DNA replication arrest events, leading to incomplete fragment amplification (*21*), or from qPCR-related variations in the gene-specific primer design and efficiency. Since qPCR amplifies only ∼200-bp regions and the terminal origins of replication were not targeted, we also recovered DNA from liposome samples by PCR followed by agarose gel analysis of the amplification products. The entire *DNArep-PLsyn* genome (within the resolution of agarose gel electrophoresis) could be recovered from diluted liposome samples (fig. S6 and fig. S7). Shorter DNA species were also observed (fig. S6 and fig. S7), suggesting that the *DNArep-PLsyn* genome may have experienced incomplete self-replication or that smaller DNA fragments were generated during PCR recovery.

Next, we sought to directly demonstrate the production of PS and intermediate enzymatic products of the reconstituted phospholipid biosynthesis pathway by liquid chromatography-mass spectrometry (LC-MS). We found that DOPS was produced, although not in high concentrations (Fig. 2G and fig. S8). Moreover, DOPA was accumulated, suggesting that CdsA may be limiting the yield of PS production. Considering that dioleoyl-phosphatidylglycerol (DOPG) accounts for 12% of the total lipids, we estimated that synthesized DOPS would represent 0.7% of the total lipid content after 16 hours incubation at 34 °C. However, it is relevant to note that LC-MS gives ensemble measurements, the obtained concentration values reflecting the average activity of all the liposomes in the sample. Individual vesicles may contain none or higher-than-average amounts of DOPS (see next section).

### High-content imaging of DNArep and PLsyn phenotypes

Having established the successful integration of the DNArep and PLsyn modules, we aimed to directly visualize the different liposome phenotypes, allowing for a more accurate classification based on activity levels. In particular, we asked whether liposome size, lamellarity, or morphology could affect or be affected by module activity. We combined fluorescence confocal microscopy with an in-house developed software called SMELDit to enable automated liposome identification, feature analysis, and image recovery from scattered data plots (see Methods). We expressed *DNArep-PLsyn* in liposomes at 34 °C and used the dsGreen and LactC2-mCherry signals as fluorescent markers for DNArep and PLsyn activity, respectively (Fig. 3A). We observed that the addition of the substrates and cofactors, and the expression of *DNArep-PLsyn* did not affect liposome sample quality (Fig. 3A, fig. S9, and movie S1). Notably, images unraveled phenotypic traits that could not be inferred from flow cytometry data, such as the presence of bright dsGreen spots in the vesicle lumen, which result from active DNA replication. We had already observed a similar phenotype during amplification of a shorter DNA self-replicator, which was attributed to an induced condensation of highly concentrated DNA (*7,22*). Here, it is interesting to see that such a phenomenon is also possible with a 3-fold longer DNA template (∼9.6 kbp vs. ∼3.2 kbp) containing more expressed genes (6 vs. 2).

**Fig. 3.**
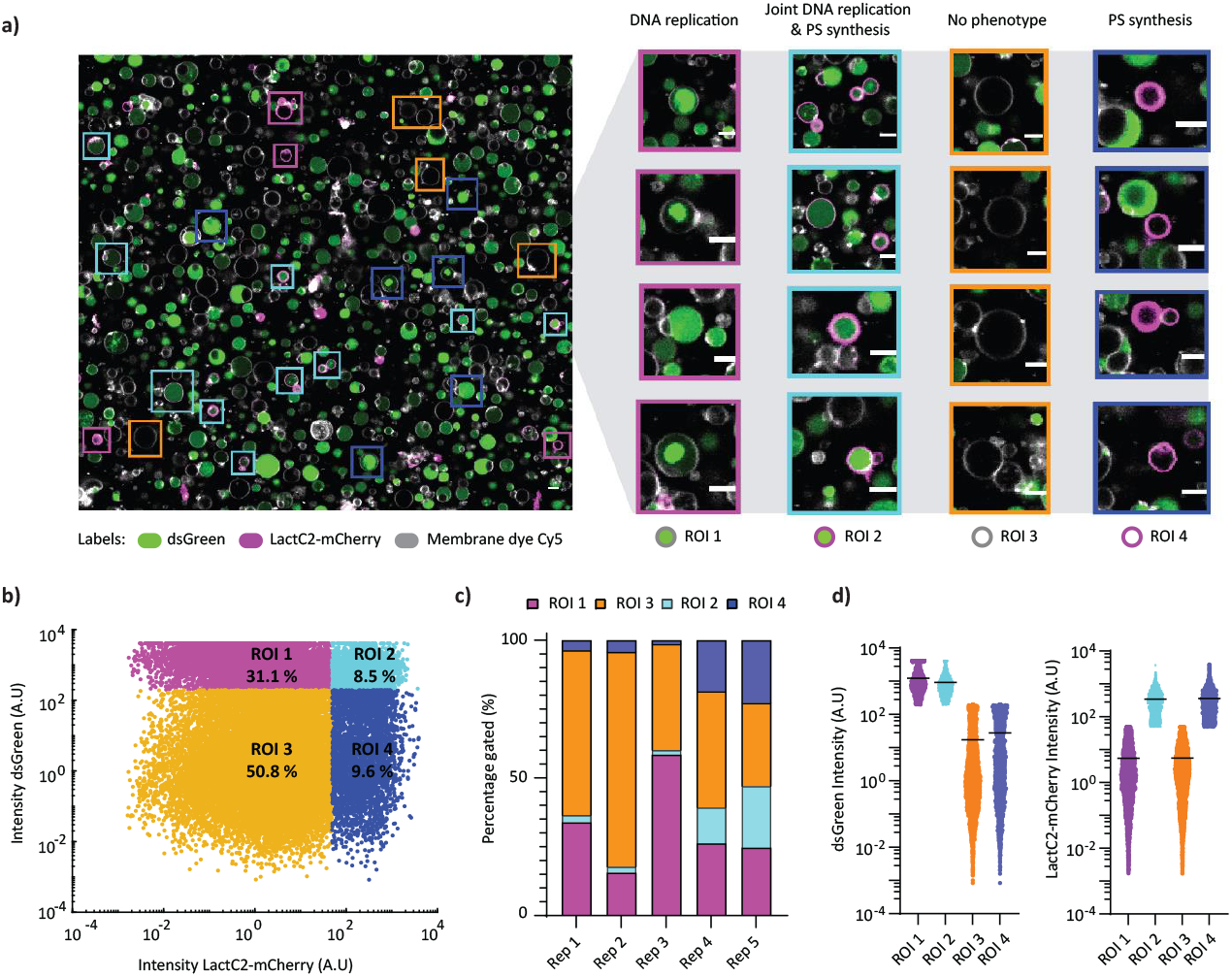
High-content imaging of DNArep and PLsyn active phenotypes. (**A**) Confocal microscopy images of gene-expressing liposomes with complete DNArep and PLsyn reaction conditions. Membrane dye (Cy5) is coloured in white, LactC2-mCherry in magenta, and dsGreen in green. Scale bar is 5 µm. Four distinct liposome phenotypes used for classification are highlighted: DNArep (ROI 1), dual DNArep and PLsyn (ROI 2), no module activity detected (ROI 3), and PLsyn (ROI 4). (**B**) SMELDit image analysis on all biological repeats (∼34,000 liposomes) builds a LactC2-mCherry vs. dsGreen phenotype map based on fluorescence intensity. Population subsets are gated into ROI 1-4 depending on the probe intensity, and are colored as in panel A. Percentages of liposomes per ROI are appended and were calculated from the pooled dataset. Phenotype maps from individual biological repeats, as well as minus DNA negative control samples can be found in fig S11. (**C**) Phenotype map (gated ROIs) from individual biological repeats. Dual phenotype region (ROI 2) is present in all replicates with at least ∼100 identified liposomes. Specifically, 273 liposomes on Rep 1, 159 liposomes for Rep 2, 94 liposomes for Rep 3, 234 liposomes for Rep 4, and 2,204 liposomes on Rep 5. (**D**) Fluorescence intensity profiles from individual liposomes across all ROIs (panel B) suggest that DNArep activity remains unaffected when coupled with PLsyn activity (left graph), and vice-versa (right graph). Each dot represents a SMELDit-identified liposome. Horizontal line indicates the mean of each data cluster.

When aggregating data from all biological replicates, over 34,000 liposomes were recognized. We generated a phenotype map corresponding to the two-dimensional plot of LactC2-mCherry vs. dsGreen signals from single vesicles (Fig. 3B). Liposomes were classified according to four different phenotypes based on intensity thresholding, akin to flow cytometry data analysis (ROI 1-4) (Fig. 2D). We found that ∼8% of liposomes, corresponding to over 2,900 liposomes, had coexisting DNA replication and DOPS synthesis (ROI 2). Vesicles with either active PLsyn (∼10%, ROI 4) or active DNArep (∼31%, ROI 1) module were more abundant (Fig. 3B). We then questioned whether liposome sizes varied across the four regions, for example as a result of membrane synthesis. Vesicle size distribution was computed for each phenotypic region (fig. S10). We observed no marked differences in the median values of the apparent diameter between active (ROI 2 and 4) and inactive (ROI 1 and 3) PLsyn module (3.7 ± 2.4 µm median across all ROIs), indicating that the yield of newly synthesized lipids is not sufficient for detectable physical growth of liposomes.

To account for the variability across biological replicates, we constructed the phenotype map for each replicate sample (Fig. 3C and fig. S11). Despite clear variations in the percentages of gated liposomes in each region, all replicates contained vesicles exhibiting simultaneous DNArep and PLsyn activity (Fig. 3C). Finally, we examined the LactC2-mCherry and dsGreen intensity values for every liposome as this may reveal differences in the efficacy of a given module when operating alone or together. From both pooled data and individual replicates, we observed no strong differences in the intensity pattern of the module activity reporter dyes between ROIs (Fig. 3D). This result suggests that DNArep activity is not lessened when coupled with PLsyn activity, and vice versa. These findings point to a robust compatibility between the two functions. In addition, some liposomes exhibit an intensity of the DOPS probe that can be over one order of magnitude higher than the average value (Fig. 3D), indicating that synthesized DOPS could represent up to 7% (0.7 × 10) of the total lipid content.

### Metabolic and activity crosstalk between DNArep and PLsyn modules

To better understand the influence that active DNArep or PLsyn modules may have on each other, we assayed liposomes expressing the full synthetic genome, this time by adding either of the two sets of substrates/cofactors (DNArep or PLsyn) (Fig. 4A). An additional condition was tested, where all DNArep substrates/cofactors were supplied, except for dNTPs. This switches DNArep module OFF but allows to study the effect of the other molecules (i.e., SSB, DSB, ammonium sulphate). We reasoned that possible inhibitory effects may arise by the substrates themselves, intermediate reaction products (e.g., lysophosphatidic acid, DOPA), or byproducts (e.g., Coenzyme A, deoxynucleoside monophosphate). Moreover, we hypothesized that DNA processing by the Φ29 DNA polymerase may either have a beneficial effect on PS synthesis by increasing the yield of synthesized enzymes through genome amplification (*19*) or have an adverse effect by hindering gene expression through collision events between DNA-interacting proteins (DSB or DNA polymerase vs. RNA polymerase) (*21*).

**Fig. 4.**
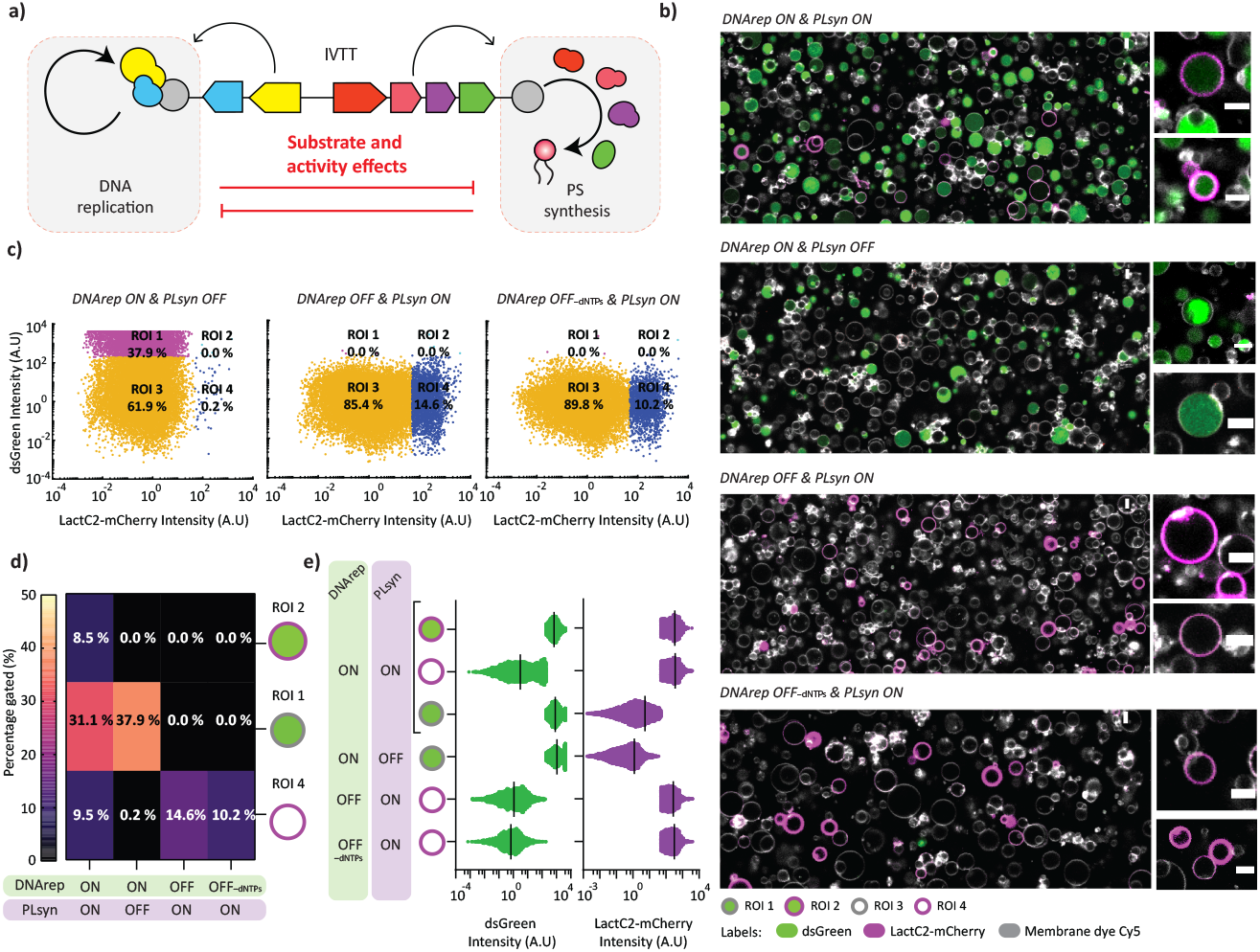
Effects of turning a module ON and OFF on the activity of the other module. (**A**) Schematic of co-active DNArep and PLsyn with an emphasis on metabolic and activity crosstalk effects. (**B**) Confocal microscopy images of liposome samples show that different substrate additions trigger a specific module activity. ON and OFF labelling indicates presence (ON) or absence (OFF) of substrates/cofactors for either DNArep or PLsyn. Liposome membrane dye (Cy5) is colored in white, LactC2-mCherry in magenta, and dsGreen in green. Scale bar is 5 µm. (**C**) Phenotype scatter plots from SMELDit image analysis (LactC2-mCherry vs. dsGreen) on all biological repeats (n = 3) show only one active module if substrates are omitted for the other one (ON or OFF state). Classified liposome subpopulations are labelled as ROI 1-4 and gated in different colors as in Fig. 3B. Appended percentages are calculated from the pooled dataset including all biological repeats. Scatter plots from the individual repeats can be found in fig. S12. (**D**) Phenotype heatmap with gated percentage values for ROIs 1,2,4 calculated across all replicates with active and/or inactive modules indicates minor crosstalk between the activity of the DNArep and PLsyn modules. Percentages for individual repeats can be found in fig S12. (**E**) dsGreen and LactC2-mCherry intensity profiles across gated ROIs are similar under both single or joint-module activity. Each dot represents a SMELDit identified liposome. Vertical lines indicate the mean value of each data cluster.

Following the same protocol as described above, we verified that DNArep and PLsyn were only active when their corresponding substrates/cofactors were present (Fig. 4, B to C, and fig. S12), confirming that nonspecific staining with dsGreen and LactC2-mCherry was negligible. Using data pooled from all biological replicates, we found that the occurrence of DNArep-active liposomes (ROI 1+2) decreased only from ∼38% to ∼31% when PLsyn substrates were supplied, while the occurrence of PLsyn-active liposomes (ROI 2+4) reduced only from ∼18% to ∼15%/∼10% (with/out dNTPs) when DNArep substrates/cofactors were supplemented (Fig. 4D and fig. S13). By examining individual replicates, we found a higher variability on the occurrence of PLsyn-active liposomes when reactions contained all DNA replication substrates/cofactors (both modules ON) compared to in their absence (fig. S13), but its cause remains to be explained. Moreover, the intensity distributions of DNArep and PLsyn activity reporters were similar regardless of the presence or absence of the substrates from the other module (Fig. 4E). Furthermore, DNA replication efficiency was similar with or without the substrates for PLsyn (Fig. 2F and fig. S14). Overall, we conclude that functional integration of the DNArep and PLsyn modules is minimally affected by metabolic crosstalk or by module co-activity.

### Influence of the genetic context on module activity

Next, we investigated whether the genetic background could influence the activity of a module. For this, we compared liposome populations with *DNArep-PLsyn* genome against liposomes with DNA templates carrying only the genes of a single module, i.e., either *DNArep* or *PLsyn*, in the presence of the full set of substrates and cofactors (Fig. 5A). We hypothesized that module activity from the *DNArep-PLsyn* genome may be compromised by sharing of resources/machinery allocated to gene expression (*23*), by impaired replication caused by strand switching of polymerizing DNAP, or by collision events between DNA interacting proteins (DNA and RNA polymerases) (*21*). All three effects would become more prominent as the number of genes increases. Alternatively, genome amplification may boost lipid biosynthesis by increasing the concentration of PLsyn enzymes (*19*).

**Fig. 5.**
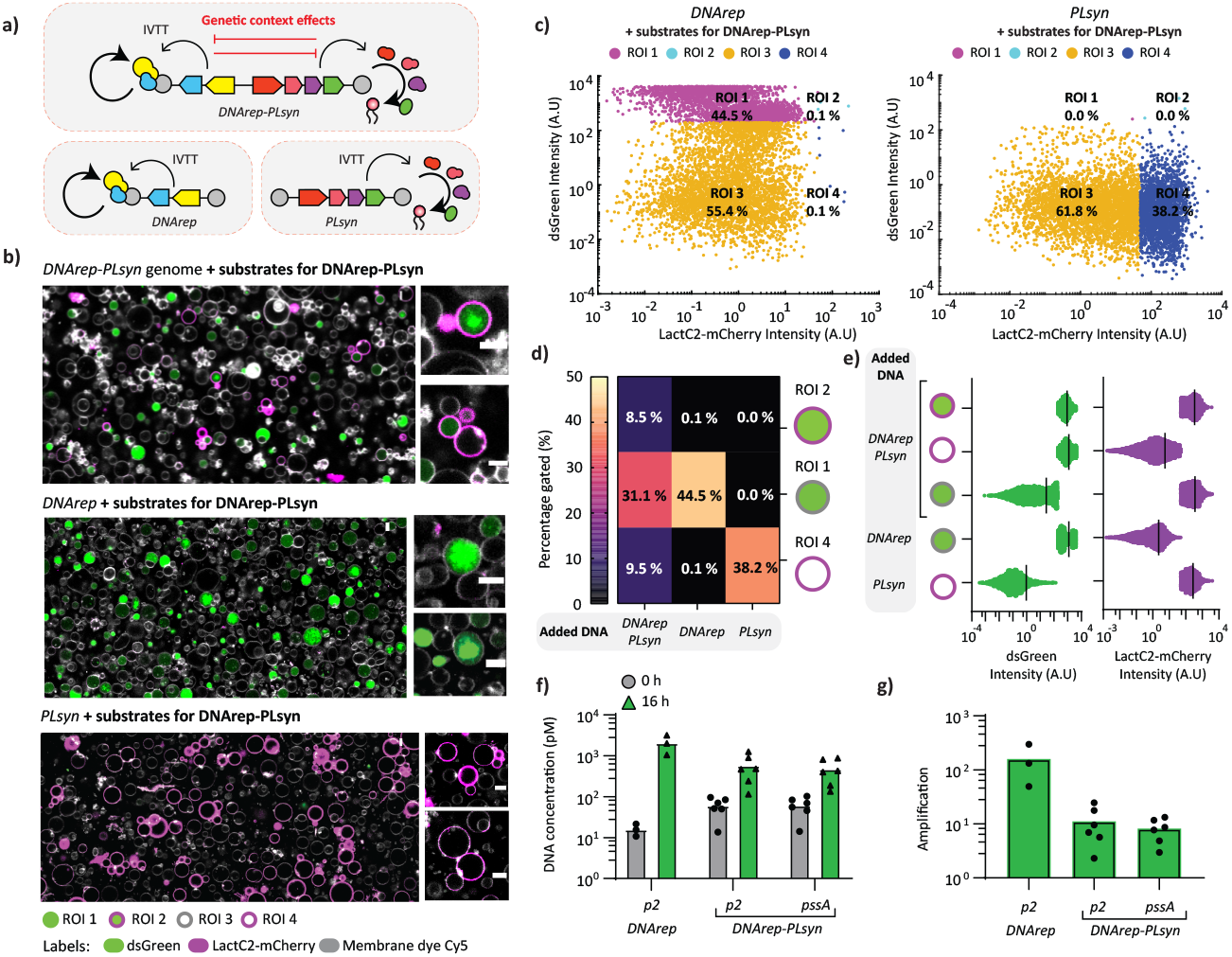
Effects of DNA template and co-expression of genetic modules on DNArep and PLsyn activity. (**A**) Schematic of the expression of the coupled (top) and separate (bottom) genetic modules from specific DNA templates. The comparison leverages the influence of genetic context on module activity. (**B**) Confocal microscopy images of liposome samples with all substrates and cofactors show DNA-specific phenotypic outputs. The used templates are indicated. Liposome membrane dye (Cy5) is colored in white, LactC2-mCherry in magenta, and dsGreen in green. Scale bar is 5 µm. (**C**) Phenotype scatter plots from SMELDit image analysis (LactC2-mCherry vs. dsGreen) on all biological repeats show only one phenotype output for each DNA program, *DNArep* or *PLsyn*. Classified liposomes in ROI 1-4 are gated in different colors. Appended percentages are calculated from the pooled data of all biological repeats. Scatter plots from individual biological replicates can be found in fig. S15. (**D**) Phenotype heatmap constructed from all repeats with different template conditions: *DNArep-PLsyn, DNArep* and *PLsyn* DNAs. (**E**) dsGreen and LactC2-mCherry intensity profiles across all ROIs have similar distributions. Each dot represents a SMELDit identified liposome. Vertical lines indicate the mean of each data cluster. (**F**) Absolute DNA quantification from liposome samples show higher DNA replication yields for the minimal self-replicator *DNArep* when compared with the *DNArep-PLsyn* genome. The targeted regions on the *pssA* and *p2* genes are indicated. (**G**) Amplification fold of *DNArep-PLsyn* and *DNArep* DNA templates calculated from qPCR data in panel F: end-point (16 hours) DNA concentration / DNA concentration at time zero. Data points represent biological repeats and bar height the mean value.

As expected, microscopy images showed that the appearance of liposome phenotypes was directed by the encapsulated DNA program (Fig. 5B to C, and fig. S15). Interestingly, the percentages of DNArep-positive liposomes were similar with and without co-expression of the *PLsyn* genes, decreasing only from ∼45% (ROI 1) to ∼40% (ROI 1+2) when *PLsyn* was co-expressed (Fig. 5D). Conversely, the percentages of PLsyn-positive liposomes dropped from ∼38% (ROI 4) to ∼18% (ROI 4+2) when *DNArep* was co-expressed (Fig. 5D and fig. S16), suggesting that PLsyn activity is more sensitive to genetic background and expression burden than DNArep activity. This effect may also limit dual-module activity in liposomes containing *DNArep-PLsyn*, explaining the higher prevalence of a single phenotype in PLsyn- and DNArep-containing liposomes (ROI 4, ∼38% on PLsyn and ∼44% on DNArep), compared to those with DNArep-PLsyn displaying both phenotypes (ROI 2, ∼8%) (Fig. 5D).

While the occurrence of liposomes exhibiting an active PLsyn module was influenced by the co-expression of *DNArep*, we noticed that the intensity distributions reporting the levels of DNArep and PLsyn activity were similar under single- and double-genetic module expression conditions (Fig. 5E). For PLsyn, this suggests that, above a detectable activity threshold, DOPS production yield was not affected by co-expression of *DNArep* genes. For DNA replication, however, dsGreen signal intensity is proportional to DNA quantity, which accounts for DNA length and amplification fold. Therefore, similar dsGreen intensities from replicated *DNArep-PLsyn* and *DNArep* templates suggest that the amplification fold of *DNArep-PLsyn* is lower than that of *DNArep* given its larger size (∼9.6 kb vs. ∼2.3 kb). To test this hypothesis, we performed absolute DNA quantitation by qPCR, confirming that *DNArep-PLsyn* replicates at a lower yield than *DNArep* (∼10-fold vs. ∼100-fold) (Fig. 5, F to G). Considering that the yield of synthesized DNAP and TP does not differ much from the *DNArep-PLsyn* or *DNArep* templates (in the absence of module-specific substrates and cofactors) (Fig. 1E and fig. S3), we speculate that the processivity of or polymerization by DNAP – and not replication initiation – might be the amplification bottleneck, especially under transcribing conditions. This hypothesis is supported by previous observations that up to a length of 6 kb the rate limiting step is initiation; over 6 kb, the DNA length becomes rate limiting (*24*).

We next examined how phenotype appearance developed in the course of gene expression. Flow cytometry data show that the highest percentage of liposomes with joint-phenotypes was reached at a later time compared to liposomes containing the genes of a single module (8 hours vs. 4 hours) (fig. S17, fig S18, and fig. S19). Time course analysis of DNA replication by qPCR showed that the maximum amplification fold was reached after 4 hours for both the *DNArep-PLsyn* and *DNArep* templates (fig. S19). This mostly reflects the DNA replication kinetics in the larger population of PLsyn-inactive liposomes expressing the full genome (ROI 1). These results demonstrate that some genetic factors slow down the dynamics of template replication when the PLsyn module is concurrently active.

### Setting up the stage for integrative evolution

Finally, we envisioned that module performance and integration could be enhanced through directed evolution. Evolving *DNArep-PLsyn* for increased functionality, e.g., a higher yield of synthesized phospholipids or faster appearance of the combined modules, would require a recursive cycle of genome library encapsulation and expression – phenotype interrogation – sorting of liposomes with the desired features – DNA recovery and amplification. We hereby streamlined the key experimental steps that are required for laboratory evolution (fig S20). First, to facilitate handling of DNA across the different stages, we here utilized the yeast-assembled plasmid as a precursor of the linear *DNArep-PLsyn* template. A cloned and sequence-verified plasmid provides a more stable template for the fast production of DNA libraries. Second, to establish a tight coupling between genotype and phenotype, we reduced the concentration of *DNArep-PLsyn* genome from 500 pM to 50 pM, which corresponds to an expected average copy number of DNA per liposome equals to one (*19*). Under these conditions a significant fraction of liposomes exhibiting combined module activation was still detected (fig. S20). Next, we screened liposomes and sorted those identified in ROI 2 by fluorescence activated cell-sorting (FACS) (∼3,000 events). The full-length genome was successfully recovered and amplified by PCR (fig. S20), and could serve as a template to start a new round. These data validate that an entire cycle of evolutionary engineering is feasible for improvement of integrated biological functions within a DNA-driven synthetic cell model.

## Conclusions

This work shows how genetically encoded functions can be integrated in a synthetic cell model. Self-replication of a DNA genome and enzymatic phospholipid synthesis were driven by a minimal transcription-translation system emulating the logic of cellular life. Although we routinely obtained more than hundred vesicles with coupled module activities per sample, joint-phenotypes did not dominate the liposome population, suggesting opportunities for improvement. Besides, challenges remain to achieve physical growth of liposomes upon lipid biosynthesis.

We found that the DNA replication and DOPS synthesis processes are fully compatible; they can simultaneously be operated without interfering with each other. However, insertion of a second genetic module reduced the occurrence of liposomes with DOPS production or the yield of amplified DNA compared to the situation in which a single genetic module was present. To alleviate this genetic burden, different designs of the *DNArep-PLsyn* genome could be tested to optimize the metabolic balance and resource allocation for gene expression. For instance, gene organization in the form of operons (*25*), or a different combination of regulatory elements (*26*–*28*) could be attempted. Considering that translation is a gene-expression bottleneck (*29*), ribosome binding sites (RBSs) of different strengths could be scanned for achieving balanced expression of the DNArep and PLsyn machineries (*30,31*). Stringent temporal control over gene expression may also be realized by implementing genetic circuits, for example ON/OFF switches regulated by specific signals (*32,33*). With this, the processes of genome replication and membrane synthesis could be separated in time, reducing competition for resources and possible clashes between DNA processing enzymes. Lastly, protein properties could be ameliorated through engineering by mutagenizing the DNA coding sequence. For example, encoding a Φ29 DNAP with higher processivity may increase the replication yield of long genomes (*34*).

While some of these modifications can be realized by rational design, we also propose to use directed evolution as an engineering tool to enhance synergy of the DNArep and PLsyn modules (*11*). We postulate that integration of rudimentary functions, such as DNArep and PLsyn, followed by evolutionary engineering, is a more effective *modus operandi* than optimizing the individual modules separately prior to combining them. Genetic diversification of the *DNArep-PLsyn* genome may occur through multiple rounds of template replication enabling the fixation of advantageous mutations directly inside liposomes (*15,35*). Alternatively, random or targeted mutations could be externally introduced, e.g., by PCR or recombineering methods (*36*), and the library of genome variants could be used as the DNA input to start an evolution cycle (fig. S20). Other interesting extensions of our work include the interconnection between the different subsystems (*37*), and the integration of more cellular modules (*4,5,8,38*) followed by system’s level evolution (*11*).

## Acknowledgments

The authors are thankful to Miguel de Vega and Alicia del Prado from the Centro de Biología Molecular Severo Ochoa, Madrid, for generously providing the purified SSB and DSB proteins. We also thank Adja Zoumaro-Djayoon for her support on the LC-MS lipidomic experiments and Ilja Westerlaken for her assistance in constructing *DNArep-PLsyn* genome batches.

## Author contributions

C.D. acquired funding, conceived and supervised the project. A.M.R.S., F.R.G., and C.D. designed the experiments. A.M.R.S. and F.R.G. performed the experiments and analyzed the data. M.T. developed SMELDit and contributed preliminary liposome experiments. L.S.H. designed and performed the molecular cloning of plasmid pY003, and helped with the DNA recovery and kinetics experiments. All authors discussed the results. A.M.R.S. and C.D. wrote the manuscript with input from all the authors.

## Data and materials availability

SMELDit code, plasmid sequences, raw LC-MS and flow cytometry data are available on GitHub (github.com/DanelonLab/pMAR3).

## SUPPLEMENTARY MATERIALS

Materials and Methods Figs. S1 to S20

Tables S1 to S2

References (*1–40*) Movie S1

