## Supplementary Information for "A synthetic cell with integrated DNA self-replication and membrane biosynthesis"

**The PDF file includes:**

Materials and Methods
Figs. S1 to S20
Tables S1 to S2

**Other Supplementary Materials for this manuscript include the following:**

Movie S1

### Materials and Methods

Buffers and chemicals

All buffers were made with MilliQ grade water with 18.2 MΩ resistivity (Millipore, USA). All chemicals were purchased from Sigma-Aldrich unless indicated otherwise.

DNA construct design

Plasmids G363 and G435 utilized for *DNArep-PLsyn* genome assembly were derived from previously cloned constructs encoding for PL synthesis enzymes (pGEMM7) (*2*) or for phi29 DNA replication proteins (G95) (*7*). Plasmid G555 harbouring the Φ29 ori-flanked *PLsyn* gene pathway was constructed by subcloning the four-gene *PLsyn* fragment from G363 (digested out with Xho1 and Nco1 restriction enzymes), into a Φ29 origins flanked vector (*7*), also digested with Xho1 and Nco1. PCR fragments for *DNArep-PLsyn* genome assembly were prepared from G363 with 5'-phosphorylated 491 ChD and 1302 ChD primers ($PLsyn_{frag}$), and from G435 with 5'-phosphorylated 492 ChD and 1289 ChD primers ($DNArep_{frag}$). Linear DNA fragments containing either of the two genetic modules were prepared by PCR using 5'-phosphorylated primers 491 and 492 ChD, and KOD Xtreme Hot Start DNA polymerase. The reaction solution contained (final concentrations) 1x Extreme Buffer (MERCK), 0.02 U/µL KOD DNA polymerase (MERCK), ~0.3 ng/µL DNA template, 300 nM forward and reverse primers, 0.4 mM of each dNTP, and MilliQ water up to 50 µL final volume. The thermocycler protocol was set to 94 °C for 2 min for polymerase activation, followed by 25-30 cycles of 10 s 98 °C, 20 s 60 to 68 °C, 30-60 s/kb at 68 °C depending on the length of the desired amplicon. All PCR amplicons were verified for correct DNA length by 0.7-1% agarose gel electrophoresis before PCR clean-up with QIAquick PCR purification kit (Qiagen). Whenever needed, DNA was purified directly from the agarose gel with a QIAquick Gel Extraction Kit (Qiagen). Both PCR and gel purification standard protocols were modified with a longer column drying step (~5 min at 10000 g) before DNA elution with MilliQ water. Purified DNAs were quantified by Nanodrop 2000c spectrophotometer (Isogen Life Science).

In vitro assembly of the *DNArep-PLsyn* genome

*DNArep-PLsyn* genome was constructed by overlap PCR from *DNArep* and *PLsyn* fragments with two main PCR steps: (i) plasmid overlap and full product DNA extension, and (ii) addition of 491 and 492 ChD primers and PCR amplification of the full *DNArep-PLsyn* product. 47 µL reactions were prepared with final concentrations of 1x Extreme Buffer (MERCK), 0.02 U/µL KOD DNA polymerase (MERCK), ~0.3 ng/µL DNA template, 0.4 mM of each dNTP, and MilliQ water. The thermocycler was programmed for 2 min at 94 °C for polymerase activation, followed by 5 cycles of (10 s at 98 °C, 20 s at 60 °C and 3 min and 40 s at 68 °C), and 20 cycles of (10 s at 98 °C, 7 min at 68 °C). For the last 20 cycles of the overlap PCR program, primers 491 and 492 ChD were added to the reaction to a final concentration of 300 nM each. The list of primers is reported in table S2.

In vivo assembly of *DNArep-PLsyn* in yeast

In vivo assembly in *S. cerevisiae* was used to build plasmid pY003, containing the *DNArep-PLsyn* genome. This plasmid was constructed from three DNA fragments: *DNArep*, *PLsyn*, and a fragment containing the yeast centromeric origin of replication *CEN6/ARS4* and the auxotrophic marker *URA3* for maintenance and selection in yeast. The *PLsyn* fragment was amplified from G363 with primers 41 ChDT and 58 ChDT, the *DNArep* fragment from G435 with primers 1289

ChD and 44 ChDT, and the *CEN6/ARS4-URA3* fragment from pRS316 with primers 56 ChDT and 57 ChDT. The PCR primers were designed to generate fragments with 60-bp overlapping ends for efficient in vivo assembly. The PCR reaction mix contained (final concentrations) 1x Extreme Buffer (MERCK), 0.02 U/µL KOD DNA polymerase (MERCK), ~0.2 ng/µL DNA template, 200 nM forward and reverse primers, 0.4 mM of each dNTP, and MilliQ water up to 50 µL final volume. The thermocycler protocol was set to 94 °C for 2 min for polymerase activation, followed by 35 cycles of 15 s at 98 °C, 20 s at 60 °C, and 60 s/kb at 68 °C depending on the length of the desired amplicon. PCR amplicons were analyzed by agarose gel electrophoresis and purified as described earlier. The DNA fragments were then pooled at a concentration of 100 fmol for the *CEN6/ARS4-URA3* fragment and 200 fmol for the other two DNA fragments. Transformation in *S. cerevisiae* CEN.PK2-1C was carried out using the lithium acetate/single-stranded carrier DNA/polyethylene glycol method (*39*). For selective growth, cells were plated on complete supplemental medium without uracil (CSM-URA) agar plates composed of 6.7 g/L yeast nitrogen base without amino acids (Formedium), 20 g/L glucose (Formedium), 0.77 g/L CSM-URA (MP Biomedicals), and 20 g/L agar (Euromedex). Plates were incubated at 28 °C for 2 days.

For pY003 isolation from *S. cerevisiae*, 15 mL cultures were grown in liquid CSM-URA media at 28 °C with shaking at 200 rpm (Infors AG CH-4103 Bottmingen). Cells were harvested at mid-log phase by centrifugation at 3,800 g for 5 minutes at 4 °C. Cell pellets were then washed with 20 mL of 100 mM Tris, pH 8.0, and centrifuged again at 3,800 g for 5 minutes at 4 °C. The pellets were resuspended in 60 µL of a freshly prepared solution containing 100 mM Tris (pH 8.0), and 10 mM DTT. To this, 3 µL of RNase A (10 mg/mL) were added, and the suspension was incubated at 30 °C for 10 minutes with shaking at 300 rpm (Eppendorf thermomixer comfort). After incubation, 8 µL of a solution containing 100 mM Tris (pH 8.0), 10 mM DTT, and 2 U/µL Zymolyase (20T, from Arthrobacter luteus, Amsbio) were added to the cell suspension. The mixture was incubated again at 30 °C for 30 minutes with shaking at 300 rpm. Spheroplasting efficiency was evaluated by measuring the optical density at 660 nm (Cary 100 Scan UV-Visible Spectrophotometer) of the samples before and after Zymolyase treatment, with an 70-100% reduction in OD indicating successful spheroplast formation. Finally, plasmid DNA was extracted using the EZ-10 Spin Column Plasmid DNA Miniprep Kit, following the manufacturer's instructions starting from the addition of Solution II (lysis buffer). DNA was eluted in 25 µL of prewarmed MilliQ water.

Due to low plasmid DNA concentration after extraction from *S. cerevisiae*, linear DNA fragments were obtained by nested PCR. The first PCR reaction (outer PCR) was performed using primers that annealed externally to the *oriL* and *oriR* flanks (50 ChDT and 51 ChDT). Subsequently, 1 µL of the product of the first PCR was used as the template for the second PCR reaction (inner PCR) using the 5'-phosphorylated primers 491E ChDT and 492E ChDT. The PCR reaction mix consisted of (final concentrations) 1x Extreme Buffer (MERCK), 0.02 U/µL KOD DNA polymerase (MERCK), 1 µL of template (Outer PCR: 1 µL of yeast-isolated plasmid; Inner PCR: 1 µL of the outer PCR), 300 nM forward and reverse primers (Outer PCR, 50 ChDT and 51 ChDT; Inner PCR, 491E ChDT and 492E ChDT), 0.4 mM of each dNTP, and MilliQ water up to 25 µL final volume. The thermocycler protocol was set to 94 °C for 2 min, followed by cycles of 10 s at 98 °C, 20 s at 60 °C, and 7.5 min at 68 °C (Outer PCR, 20 cycles; Inner PCR, 35 cycles). PCR amplicons were analyzed by agarose gel electrophoresis and purified as described earlier. The final *DNArep-PLsyn* amplicon was sequence-verified using

Oxford Nanopore technology (Plasmidsaurus). Three single-nucleotide substitutions were identified but these mutations did not hinder activity of the two modules.

The lists of plasmids and primers are reported in table S1 and table S2, respectively.

Purification of SSB, DSB, and LactC2-mCherry

Purified SSB and DSB Φ29 auxiliary proteins were produced and stored in –80 °C as previously described in (*7*). SSB stock concentration was 10 mg/mL, stored in a buffer with 50 mM Tris, pH 7.5, 60 mM ammonium sulphate, 1 mM EDTA, 7 mM β-mercaptoethanol (BME), and 50% glycerol. DSB stock concentration was 10 mg/mL, stored in a buffer with 50 mM Tris, pH 7.5, 0.1 M ammonium sulphate, 1 mM EDTA, 7 mM BME, and 50% glycerol. LactC2-mCherry protein stock was produced and stored in –80 °C as described in (*19*). LactC2-mCherry stock concentrations were determined by a Bradford assay, and stored in a buffer with 50 mM HEPES-KOH, pH 7.5, 150 mM NaCl, and 10% glycerol. Before liposome staining, the protein stock vial was centrifuged at maximum speed (Eppendorf Centrifuge 5415 R) for 10 min to spin down protein aggregates. If necessary, LactC2-mCherry protein stock was diluted in homemade PURE buffer (PB) consisting of 180 mM potassium glutamate monohydrate, 14 mM magnesium acetate tetrahydrate, and 20 mM HEPES at a pH of 7.6 (adjusted with potassium hydroxide) before usage.

In-liposome gene expression

Lipid-coated beads were prepared as explained in (*2,7*) with minor modifications. A lipid mixture was prepared with chloroform-dissolved lipids (Avanti Polar Lipids) in a 5-mL round-bottom flask. The solution contained 49.5% DOPC, 33.7% DOPE, 12% DOPG, 3.8% 18:1 CL, and 1% DSPE-PEG-biotin mass composition, for a total mass of lipids of 2.02 mg. For confocal microscopy experiments the primary lipid composition was adjusted to include 0.05% mass of DOPE-Cy5. The resulting mix was supplemented with 25.4 µmol of rhamnose (Sigma-Aldrich) dissolved in methanol. 600 mg of 212–300-µm glass beads (Sigma-Aldrich) were added to the 5-mL round-bottom flask containing the lipid/rhamnose solution, and the solvent was evaporated with a rotary evaporator (Heidolph) for 2 hours at room temperature and 200 mbar. The lipid-coated beads were recovered and further dried in a desiccator overnight. The dried lipid-coated beads were flushed with argon and stored at −20 °C until use. In 1.5 mL Eppendorf tubes, PURE*frex2.0* (GeneFrontier) reactions were assembled as recommended by the manufacturer using 500 pM DNA template (if not mentioned otherwise) and 0.75 U/µL of Superase·In RNase inhibitor (Thermo Fisher). DNA assembled in vitro was used in all experiments, except for those described in fig. S4 and fig. S20, where the genome assembled in yeast was employed. When indicated, the IVTT mixture was supplemented with the required substrates and cofactors for DNA replication (300 µM dNTPs, 20 mM ammonium sulphate, 0.75 mg/mL SSB, and 0.21 mg/mL DSB, final concentrations) or/and PL synthesis (1 mM CTP, 500 µM G3P, 500 µM L-Serine, and 5 mM β-mercaptoethanol, final concentrations, the oleoyl-CoA precursor was added in a next step described below). 10 mg of lipid-coated beads freshly desiccated for 20-30 min were added to 20 µL of swelling solution. The sample in Eppendorf tube was kept in a 4 °C room for gentle rotation with an automatic tube rotator (VWR) for 30 min, and was then subjected to four freeze-thaw cycles with alternating steps of dipping in liquid nitrogen and thawing on ice. Liposomes were recovered with a cut pipette tip and transferred to a PCR tube with DNase One (NEB) added to a final concentration of 0.1 U/µL. For PLsyn activity, the sample was further transferred to another PCR tube with a pre-deposited 1 µg O-CoA (Avanti

Polar Lipids) film, prepared as described in (*2*). With the added liposome suspension, the final O-CoA concentration was 176 μM, and when needed (i.e., liposome suspension volumes changed), the pre-deposited O-CoA quantity was adjusted to maintain the same final concentration. Samples were incubated at 30, 37, or 34 °C for 16 hours. A maximum of 15 μL liposome suspension was handled per PCR tube. If higher volumes were needed, the suspension was distributed across different PCR tubes with independently added O-CoA dried lipid film.

Flow cytometry

In a PCR tube, a liposome suspension (1.5 μL) was mixed in a 1:1 ratio with a 1000x diluted dsGreen (Lumiprobe) stock solution in PB and the sample was left to incubate in the dark at room temperature for 30 min. The 3 μL liposome-dsGreen mixture was further diluted by adding 197 μL of a 1:1000 dsGreen:PB solution. To remove possible remaining glass beads, the 200 μL solution was filtered through a 35-μm nylon mesh of a cell-strainer cap from 5-mL round-bottom polystyrene test tubes (Falcon). With a large volume pipette tip, 138.5 μL of the filtered solution was transferred to a 2 mL round-bottom tube to which 1.5 μL of 1:1000 dsGreen:PB solution and 10 μL of LactC2-mCherry probe were added to obtain a final LactC2-mCherry concentration of 300 nM and a final volume of 150 μL. The sample was incubated for 1 hour before injection in a FACSCelesta flow cytometer (BD Biosciences) set up with a 488-nm laser and 530/30 filter for detection of dsGreen, and with a 561-nm laser and 610/20 filter for detection of LactC2-mCherry. Photon multiplier tube voltages were 375 V for forward scatter, 260 V for side scatter, 370 V for LactC2-mCherry, and 550 V for dsGreen detection. Loader settings were set to 50 μL injection volume with no mixing and 800 μL wash between sample runs. For each sample ~20000 events were recorded. The raw data were analyzed and pre-processed using Cytobank (https://community.cytobank.org/) to filter out possible aggregates and liposome debris as previously described in (*19*). Text files with all SSC-A, dsGreen, and LactC2-mCherry intensity values were exported from Cytobank and plotted with MATLAB.

Confocal microscopy

Liposome samples (2 μL) were transferred in custom-made glass chambers functionalized with BSA-biotin:BSA and Neutravidin, as previously described in (*2*), and pre-filled with 13 μL of a staining solution (1X dsGreen and 3 μM LactC2-mCherry diluted in PB). Chambers were incubated in the dark at room temperature for 1 hour. Confocal microscopy imaging was carried out on a Nikon Eclipse Ti (NIS-Elements AR software) using a 100x oil immersion objective. Laser settings for image acquisition were set to: 488-nm laser with 20 HV, -10 offset, and 1.0 intensity for dsGreen, 561-nm laser with 50 HV, -10 offset, and 1.00 intensity for LactC2-mCherry, and 640 nm laser with 95 HV, -5 offset, and 5.00 intensity for Cy5 membrane dye. Each sample was imaged by automated acquisition of 10 by 10 fields of view, stitched together with a ~5% overlap. The sample height was adjusted manually to detect as many liposomes as possible, while also avoiding background debris.

Image analysis

Confocal images were analyzed manually with Fiji (*40*) and automatically with SMELDit, an in-house-developed MATLAB script to automatically extract single liposome features while indexing each analyzed liposome. In short, the Cy5 and mCherry channels of each image are combined and convolved with a Laplacian filter kernel to determine membrane boundaries. To set what pixels belong to the inside of each liposome a binarization step, with a consistent cutoff based on previous data, followed by a filling and erosion step were utilized. The resulting binary image displayed separate segments, each representing the lumen of individual liposomes. This step was followed by an additional selection for filtering out the segments that could correspond to lipid aggregates or other noise sources. Then, all segments were analyzed individually for a circularity ($C$) check defined by $C = \frac{P^2}{4\pi A}$, where $P$ is the perimeter length and $A$ is the area of the segment. A 'true liposome' threshold was set for $C$ values between 0.5 and 2.0. LacC2-mCherry aggregates were filtered out by rejecting events whenever LactC2-mCherry intensities were higher than a pre-set cutoff inside the lumen. The segments that passed these extra filtering steps were considered liposomes and were saved individually as 60 by 60-pixel cropped images with a given ID number. For each individual liposome SMELDit measures the apparent radius, average Cy5 intensity and variance on the membrane, average LactC2-mCherry intensity and variance on the membrane, and average dsGreen intensity and variance on the lumen. Per sample, SMELDit displays all single-liposome extracted data in a scatter/histogram interactive GUI on which the user can draw regions of interest (ROI) for extracting information about liposome subpopulations. Here, ROI 1-4 were defined in SMELDit using negative controls for thresholding. Finally, once ROIs are drawn, example liposomes from the ROI are retrieved as a liposome-montage. ROI liposome data can be saved separately and ROI coordinates can also be saved and transferred to analyze another sample. The SMELDit MATLAB code is available on GitHub (github.com/DanelonLab).

Quantitative PCR analysis

One to two microliter of liposome suspension was collected, incubated 15 min at 75 °C for DNAse I heat inactivation, and 100x diluted in MilliQ water. Ten microliter qPCR reactions were prepared with 1x PowerUP SYBR Green Master Mix (Applied Biosystems), 400 nM of each forward and reverse primer (976/977 ChD for *p2,* 980/981 ChD for *p3,* 1125/1126 ChD for *pssA,* 1119/1120 for *plsB,* 1410/1411 ChD for *plsC,* 1408/1409 ChD for *cdsA*), and 1 µL of the diluted liposome sample. Solutions were transferred to a qPCR 96-well plate (Thermo Fisher) that were sealed with an adhesive transparent film (Thermo Fisher) and spined down for 15 seconds. Measurements were performed on a Quantstudio 5 Real-Time PCR instrument (Thermo Fisher) using the protocol: 2 min at 50 °C, 5 min at 94 °C, 45 cycles of 15 sec at 94 °C, 15 sec at 56 °C, 30 sec at 68 °C, 5 min at 68 °C, and a final melting curve stage from 65 °C to 95 °C. Sample DNA concentrations were calculated from standard curves generated using DNA templates of known concentrations ranging from 1 fM to 1 nM (9 µL of qPCR reaction + 1 µL of DNA). Data were further analyzed with Quantstudio Design and Analysis software v1.4.3 Software (Thermo Fisher).

Recovery of *DNArep-PLsyn* DNA from liposomes

Liposome samples diluted 100 times in MilliQ water for qPCR measurement were utilized also for DNA PCR recovery. Reactions were set up to either amplify three fragments or a single, near full-length, fragment from the *DNArep-PLsyn* genome. Twenty to 50 µL reaction solutions were assembled in a 1x Xtreme Buffer with 300 nM of each primer (all primers and details about the corresponding PCR amplification targets can be found on table S2), 0.4 mM of each dNTP, 2-5 µL of the diluted liposome solution, and 0.02 U/µL KOD DNA polymerase. The thermal cycler was programmed to follow 2 min at 94 °C for polymerase activation, and 30 cycles of (98 °C for 10 sec, 60 °C for 20 sec, 68 °C for 2 min for fragments A,B,C, and 5 min for one-fragment PCR-recovery). PCR products were analyzed by agarose gel electrophoresis.

Bulk IVTT reactions

Ten microliter bulk IVTT reactions were performed with PURE*frex* 2.0 using 500 pM DNA template according to the manufacturer's guidelines. To visualize synthesized protein products, the reaction was supplemented with 1 µL GreenLys solution (FluoroTect GreenLys, Promega) and incubated for 16 hours at 37 °C. Samples were then supplemented with 1 µL of RNase A (4 mg/mL) and 1 µL of RNase One (10 U/µL), and incubated for 1-2 hours at 37 °C for complete RNA digestion. 10 µL of the RNA-digested sample were mixed with 1x Laemmli sample buffer and 10 mM of DTT, final concentrations. The reaction mixtures were incubated at 95 °C for 5 min and loaded on a 12% SDS-PAGE gel that was run for 1 hour at 100 V, followed by another 50-60 min at 130-160 V. GreenLys labelled proteins were visualized on a fluorescence gel imager (Typhoon, Amersham Biosciences) using Cy2 (488 nm), Cy3 (532 nm), and Cy5 (635 nm) lasers with band-pass filters of 515-535 nm for Cy2, 560-580 nm for Cy3, and 655-685 for Cy5. Laser PMT voltages were set to 500 for Cy2 and automatic adjustment for Cy3 and Cy5. The SDS-PAGE gel was further stained overnight with Instant Blue (expedeon), destained the next day with MilliQ water, and visualized with a ChemiDoc imaging system (Bio-Rad).

Analysis of lipid content by LC-MS

Two to four microliters of liposome suspension were diluted 10x in a sample preparation solution consisting of a 1:1:98 ratio of 0.5 M EDTA:200 mM acetylacetone:100% methanol. The solution was sonicated for 10 min, and centrifuged at 16.1 g and room temperature for 5-10 min. 15 µL of the supernatant with the soluble lipid phase were transferred to a 25 µL glass insert within a 2 mL LC-MS glass vial, further diluted 4x with the sample preparation solution, and stored under argon at −20 °C until use (less than a week). For LC-MS sample analysis, ~10 µL sample kept at 4 °C were injected into a 6460 Triple Quad LC-MS stocked with a ACQUITY UPLC Peptide CSH C18 Column with a mobile phase A of 0.05% ammonium hydroxide and 2 mM acetylacetone in water, and a mobile phase B of % 2-propanol, 20% acetonitrile, 0.05% ammonium hydroxide and 2 mM acetylacetone, at a flow rate of 300 µL/min and column temperature of 60 °C. An A:B ratio of 70:30 was set to equilibrate the column. Upon sample injection, the A:B ratio was slowly changed to 100% mobile phase B and kept for two min. Then, the 70:30 ratio was gradually restored and kept until the end of each sample run. Transitions were established in our previous work (*52*). Data were analyzed with Skyline-daily. Peak areas were exported and normalized to DOPG peak areas. When two injections were done per sample, the averaged peak area was considered.

**Fig. S1.**

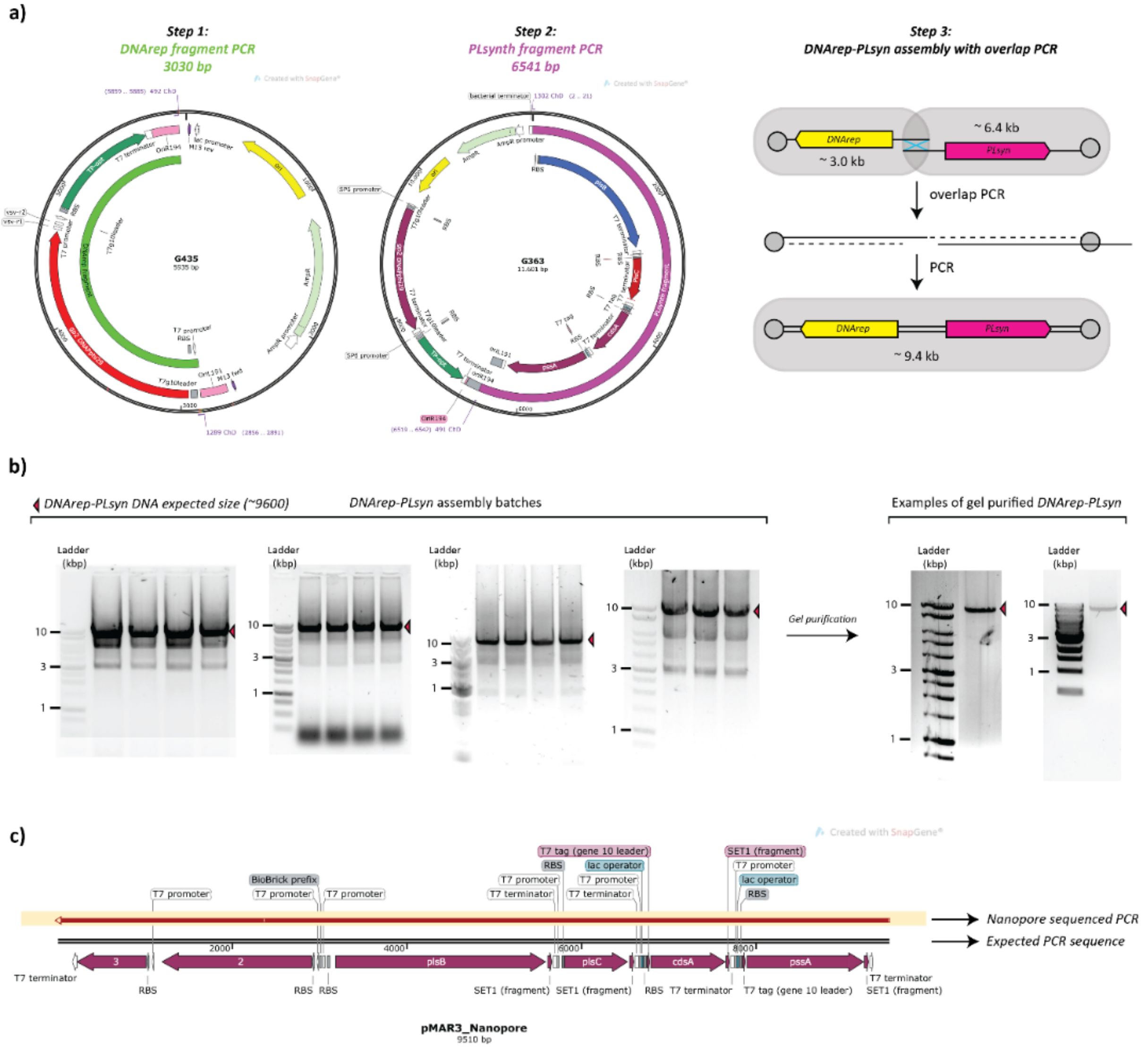


**Fig. S1.** ***DNArep-PLsyn* genome assembly by overlap PCR. a)** Experimental steps to prepare *DNArep-PLsyn* genome. Steps 1 and 2 correspond to the preparation of *DNArep* and *PLsyn* fragments by PCR from G435 and G363 plasmids (table S1). In step 3 the DNA fragments are stitched by overlap PCR to obtain the *DNArep-PLsyn* genome (~9.4 kb). **b)** Agarose gel electrophoresis of different *DNArep-PLsyn* genome assembly batches without (left) and without (right) gel purification. **c)** DNA sequence alignment (from SnapGene) between the expected *DNArep-PLsyn* sequence and a sequence-verified *DNArep-PLsyn* assembly batch (Oxford Nanopore sequencing, Plasmidsaurus).

Fig. S2.

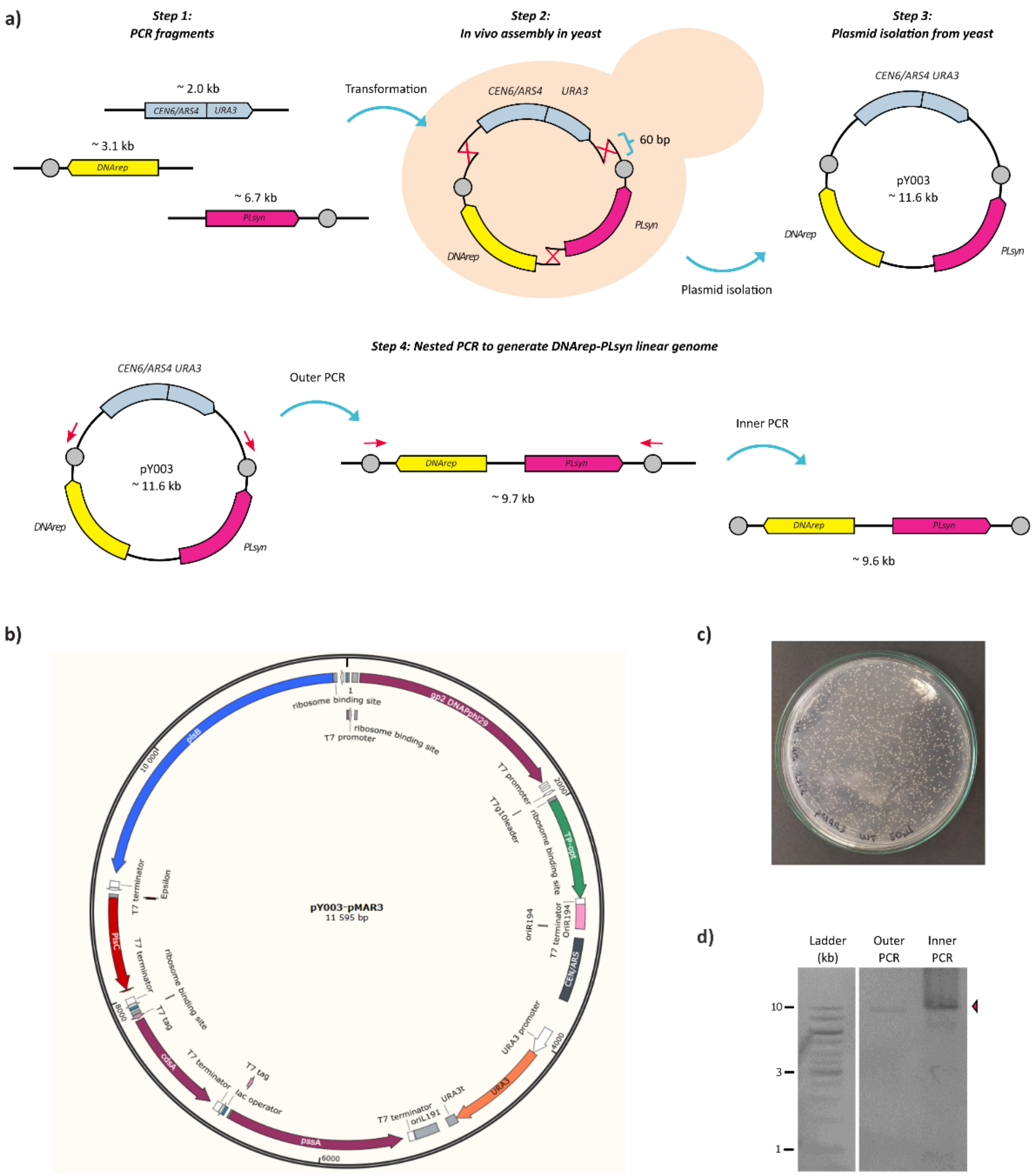


**Fig. S2.** ***DNArep-PLsyn* genome assembly in yeast. a)** Experimental steps to prepare *DNArep-PLsyn* genome. Step 1 entails the generation of the *DNArep*, *PLsyn,* and *CEN6/ARS4 URA* fragments by PCR from G435, G363, and pRS316, respectively (table S1). Step 2 involves the transformation and assembly of the DNA fragments in yeast. Fragments have 60 bp overlapping sequences for homologous recombination. Step 3 represents the extraction of the assembled plasmid, pY003 (~11.6 kb), which contains the *DNArep-PLsyn* genome. Step 4 illustrates the generation of the linear *DNArep-PLsyn* genome by nested PCR. **b)** pY003 plasmid map (from SnapGene). **c)** Transformation plate after in vivo assembly of pY003. **d)** Agarose gel electrophoresis of the nested PCR to generate the linear *DNArep-PLsyn* genome.

**Fig. S3.**

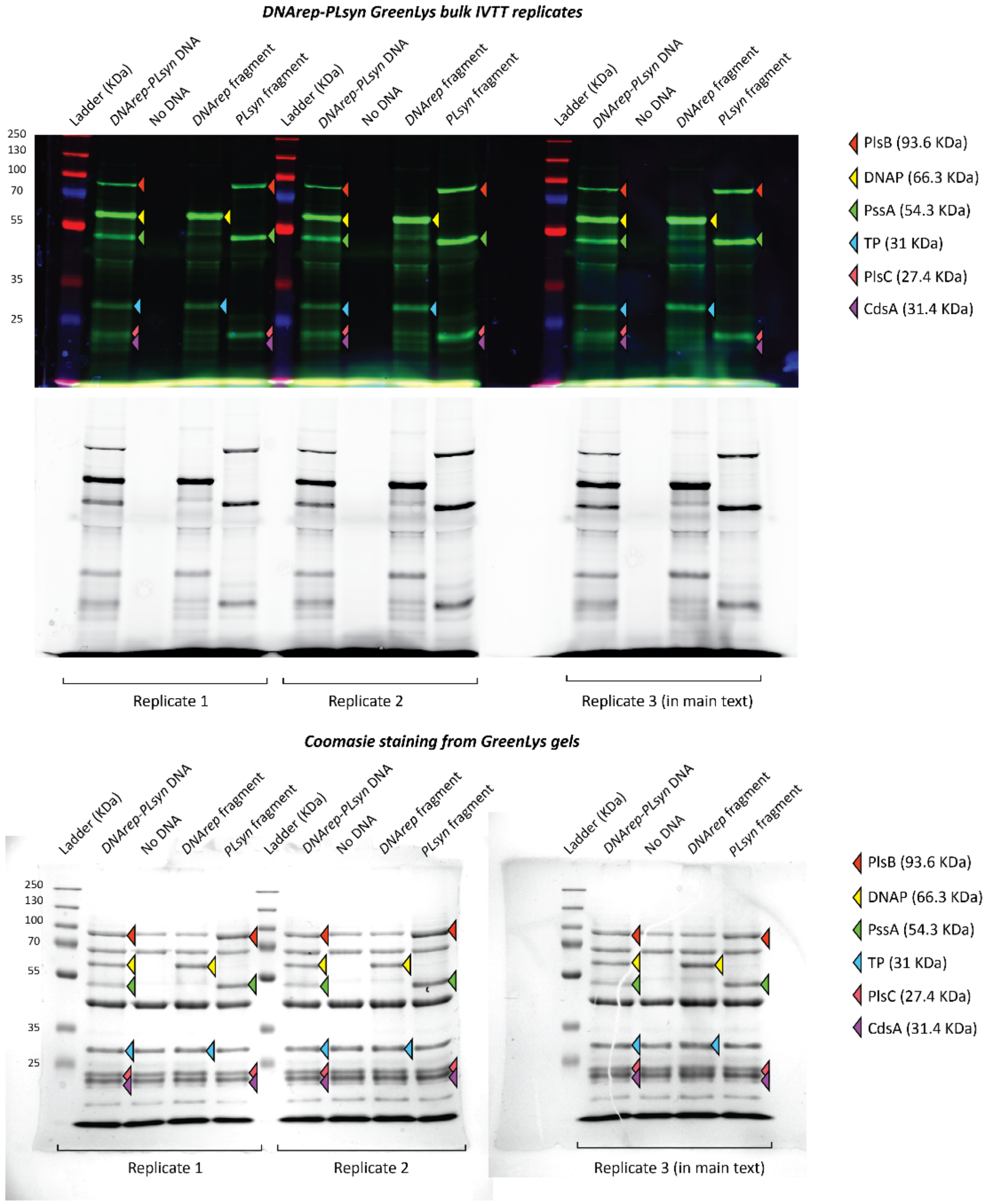


**Fig. S3.** Biological repeats of bulk IVTT protein production with *DNArep-PLsyn*, *DNArep*$_{frag}$ and *PLsyn*$_{frag}$ as DNA templates. SDS-PAGE gels with Green-Lys protein labelling (upper gel), and Coomassie protein staining (bottom gel).

Fig. S4.

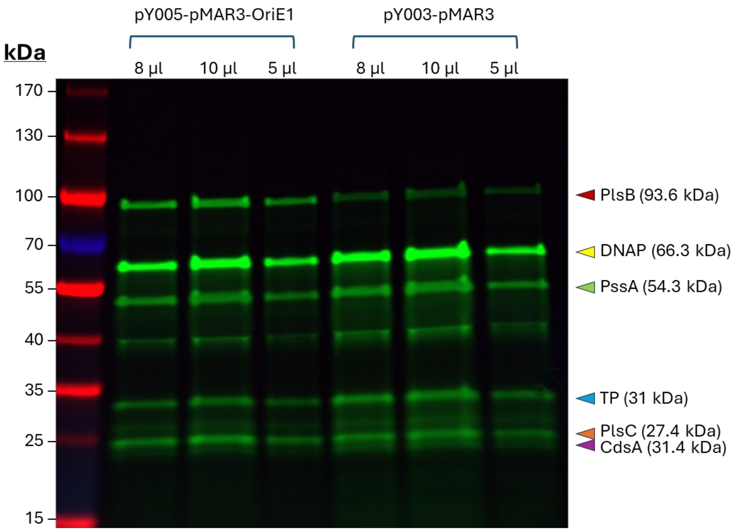


**Fig. S4. Visualization of synthesized protein from a *DNArep-PLsyn* template assembled in yeast.** SDS-PAGE gel with Green-Lys protein labelling. Protein expression patterns are shown with two versions of the template (only pY003 was used for activity assays) and different volumes of loaded samples, as indicated. DNA concentration was 2 nM and the IVTT solution was incubated for 16 hours at 37 °C.

**Fig. S5.**

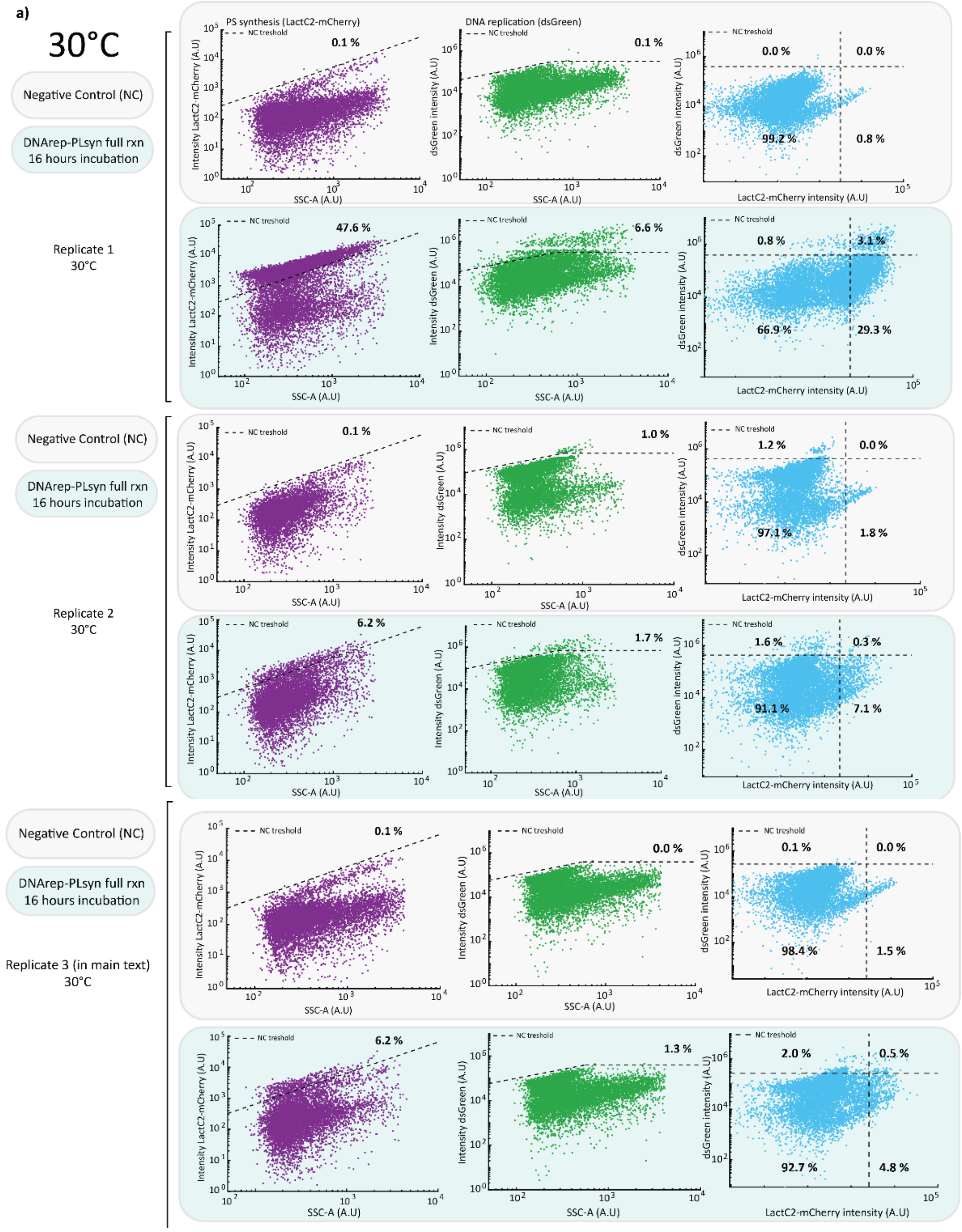


**Fig. S5** continues next page

**Fig. S5.**

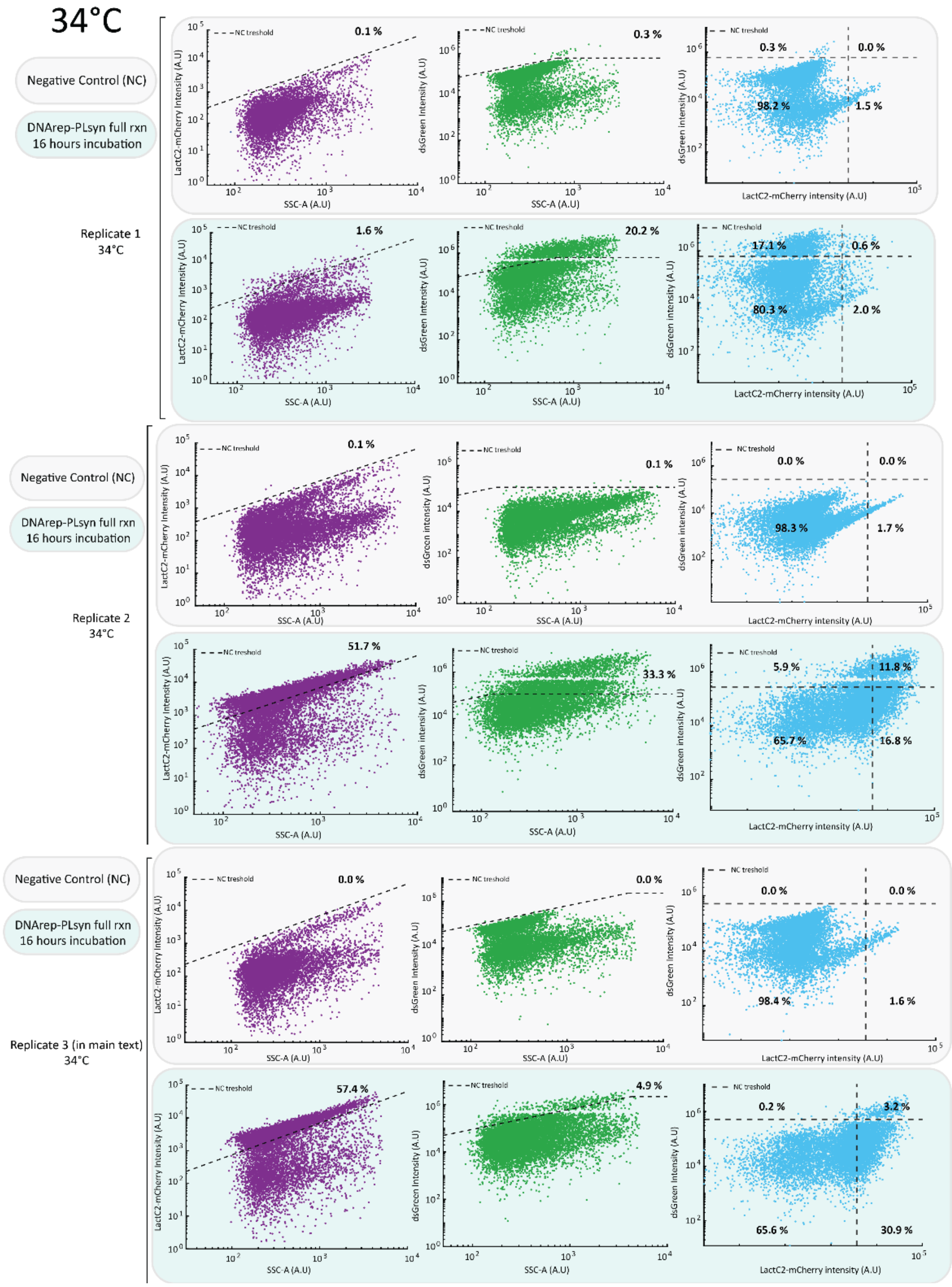


**Fig. S5** continues next page

**Fig. S5.**

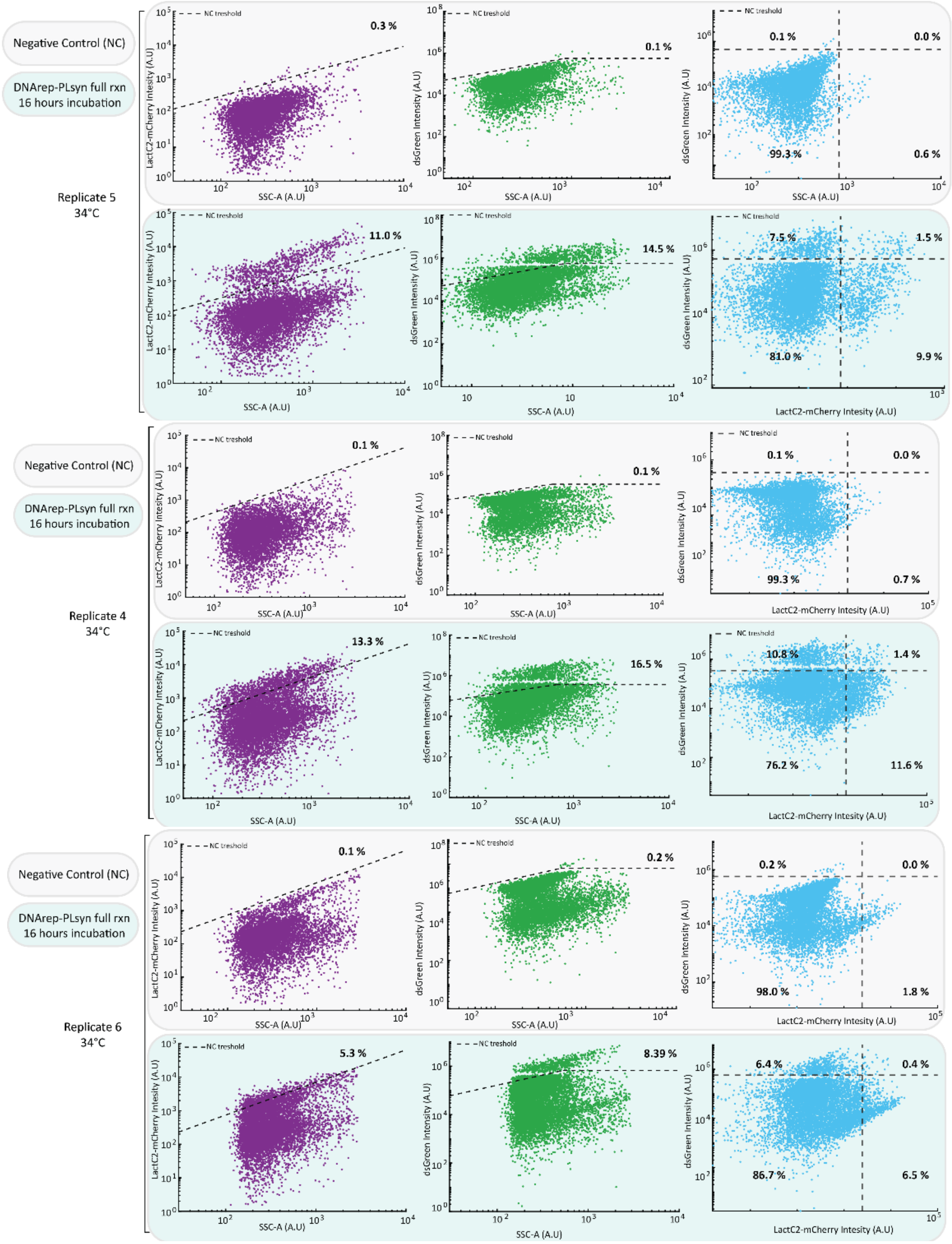


**Fig. S5** continues next page

**Fig. S5.**

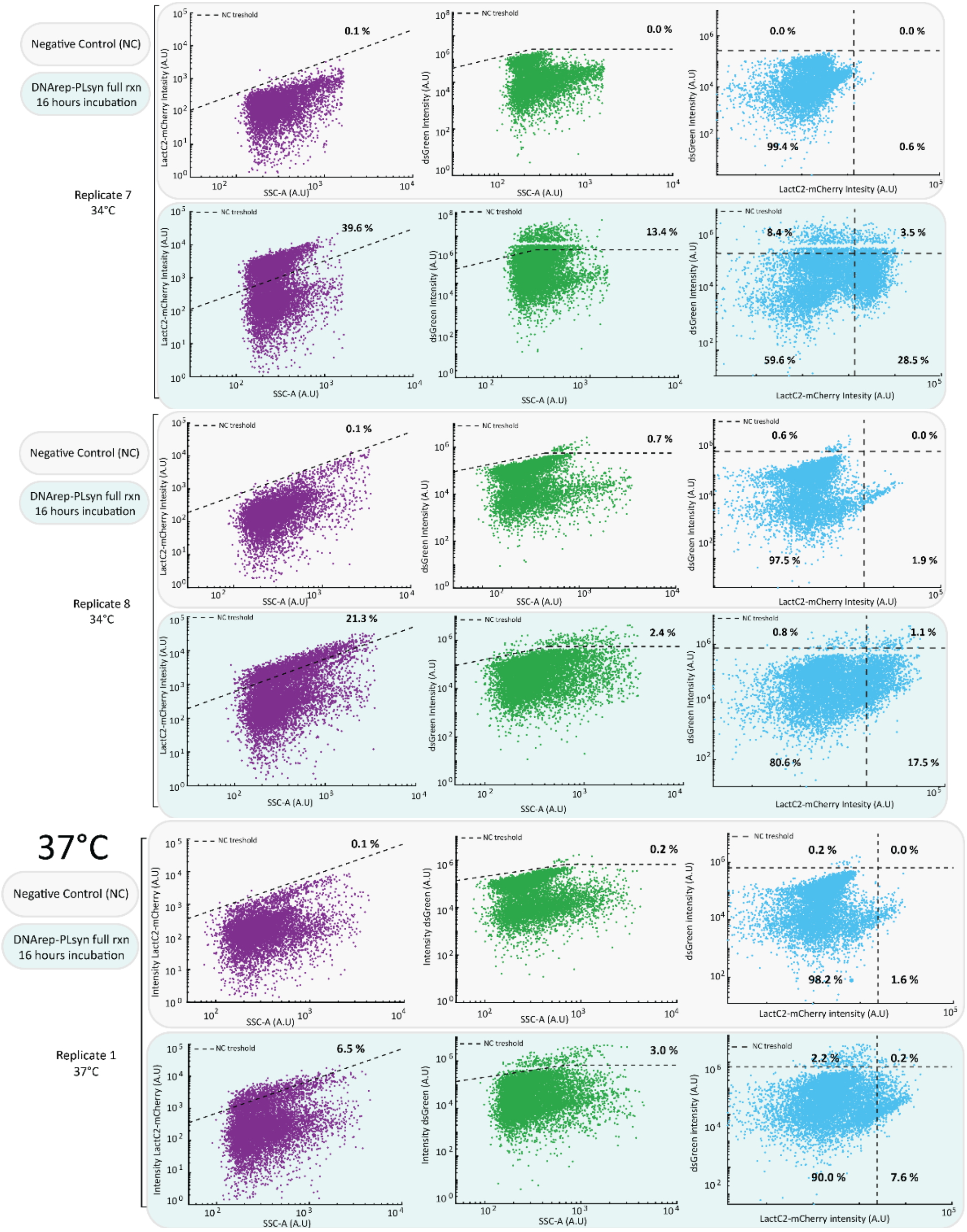


**Fig. S5** continues next page

**Fig. S5.**

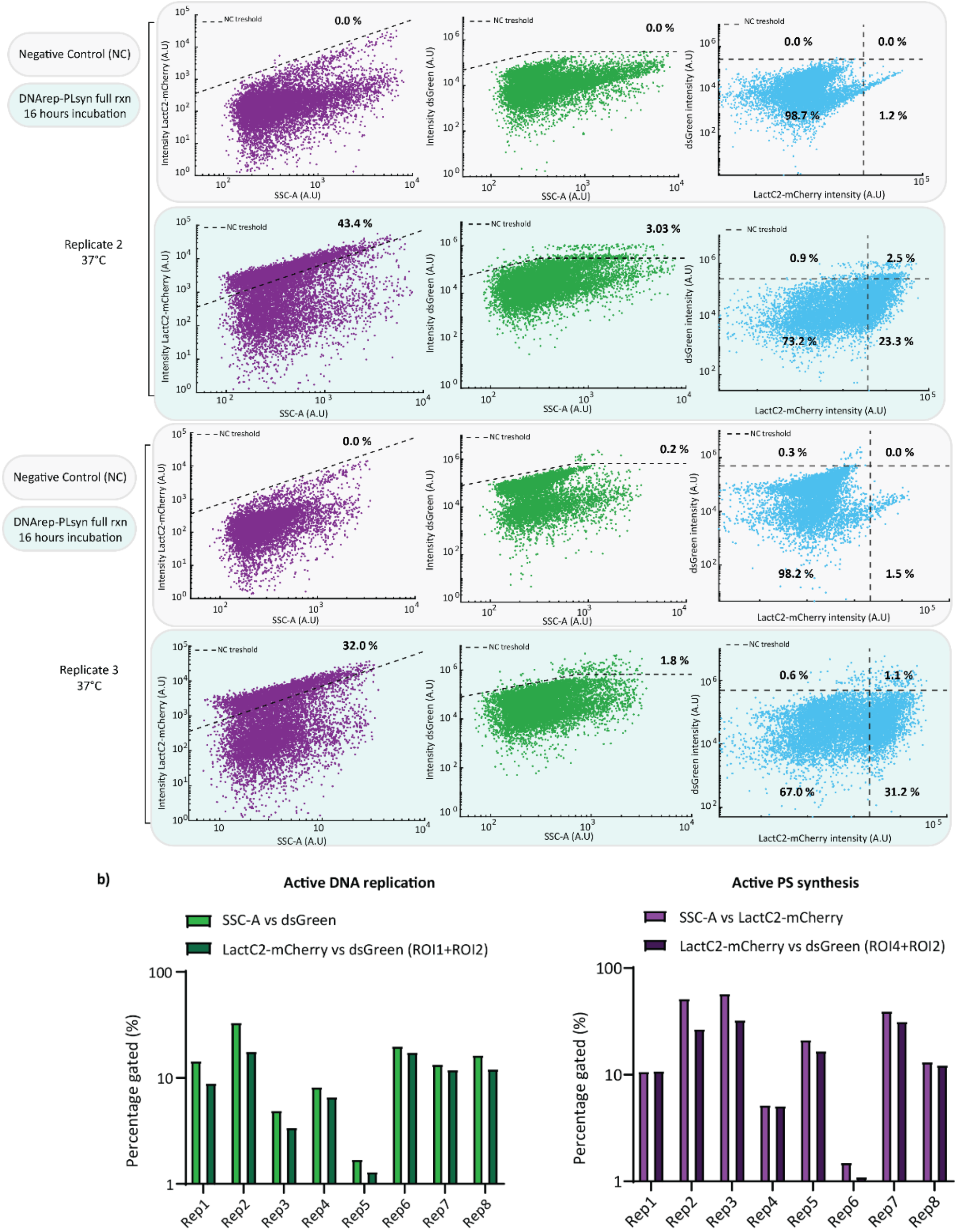


**Fig. S5. a)** Flow cytometry scatter plots from all the biological repeats of *DNArep-PLsyn* activity assays shown in Fig. 2. Samples were incubated at 30, 34, or 37 °C, as indicated. **b)** Bar plot representation of the percentage of gated liposomes calculated from the SSC-A and fluorescence signals, as specified in the legend, for 8 biological replicates. DNA-rep- and PLsyn-active liposomes were classified based on intensity thresholds as described in the Methods section.

**Fig. S6.**

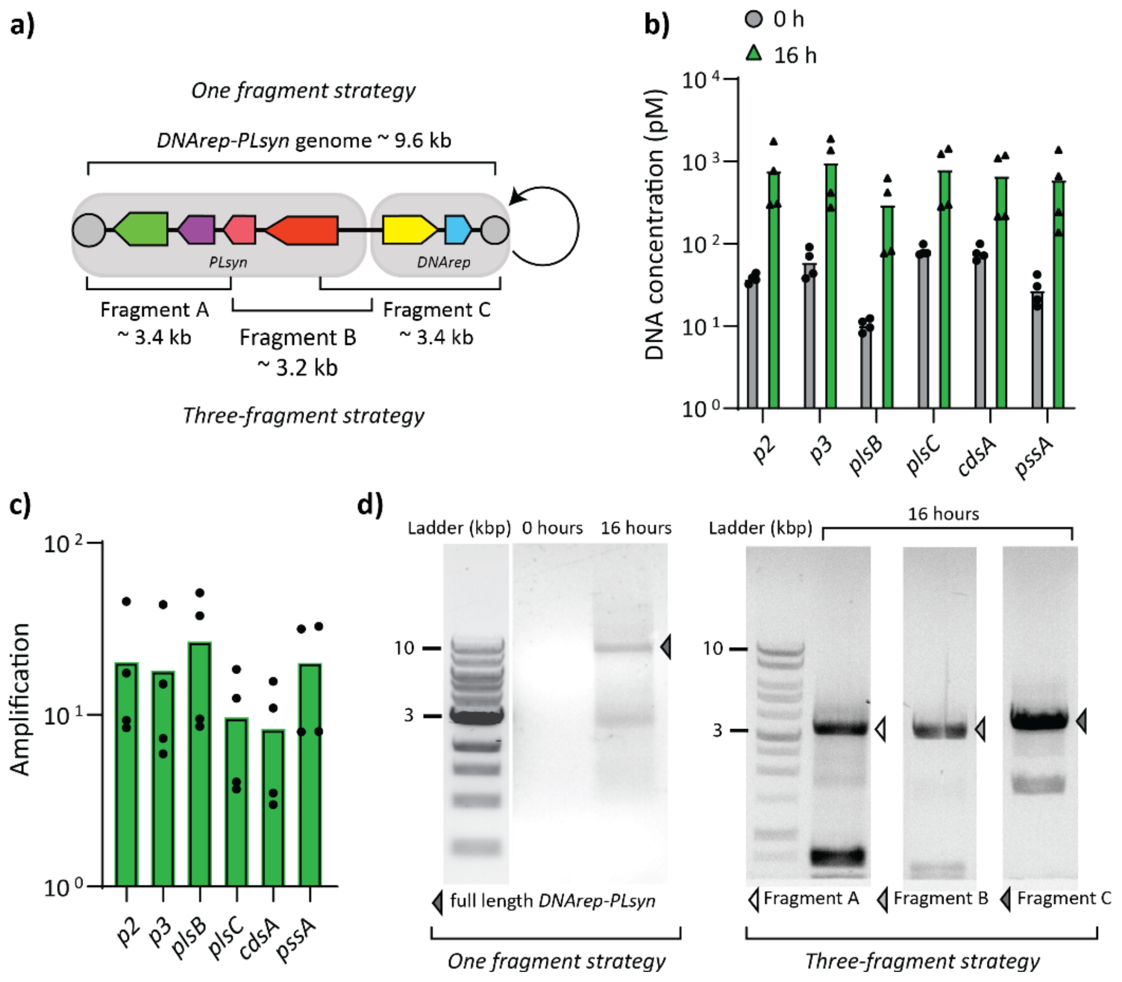


**Fig. S6. *DNArep-PLsyn* genome self-replicates in liposomes under joint-module reaction conditions at 34 °C. a)** Illustration of the two DNA recovery strategies consisting of either three or one fragment that was PCR-amplified from *DNArep-PLsyn*. **b)** Absolute DNA quantification of DNArep-PLsyn containing liposomes at time zero (dark green) and after 16 hours (light green) of sample incubation. The gene names of the targeted regions (~200 bp) are indicated. **c)** DNA amplification fold calculated from the data shown in b. Each data point within a target gene represents a biological repeat. **d)** Agarose gel electrophoresis of PCR-recovered *DNArep-PLsyn* with both PCR recovery strategies as illustrated in panel a. On the left, the ladder lane was cropped from the same gel image and appended.

**Fig. S7.**

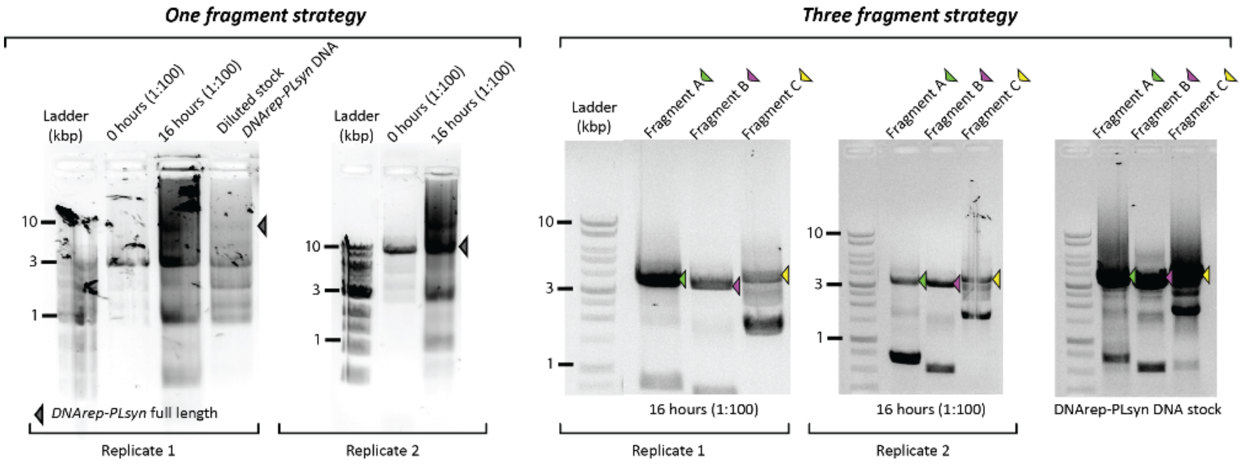


**Fig. S7.** Agarose gel electrophoresis of PCR-recovered DNA from the *DNArep-PLsyn* template isolated from liposome samples. A one- or three-fragment (A, B, C) recovery strategy was used, as indicated.

**Fig. S8.**

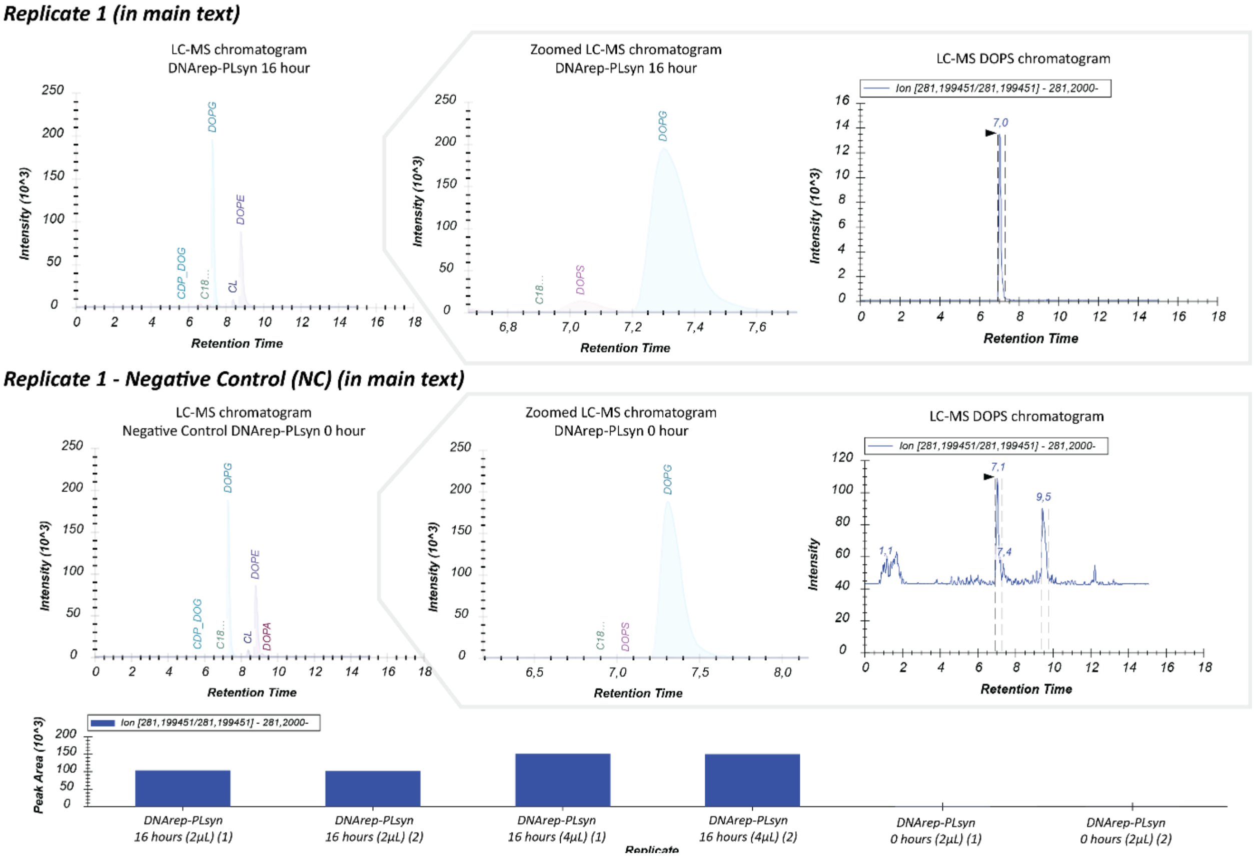


**Fig. S8** Continues next page.

**Fig. S8.**

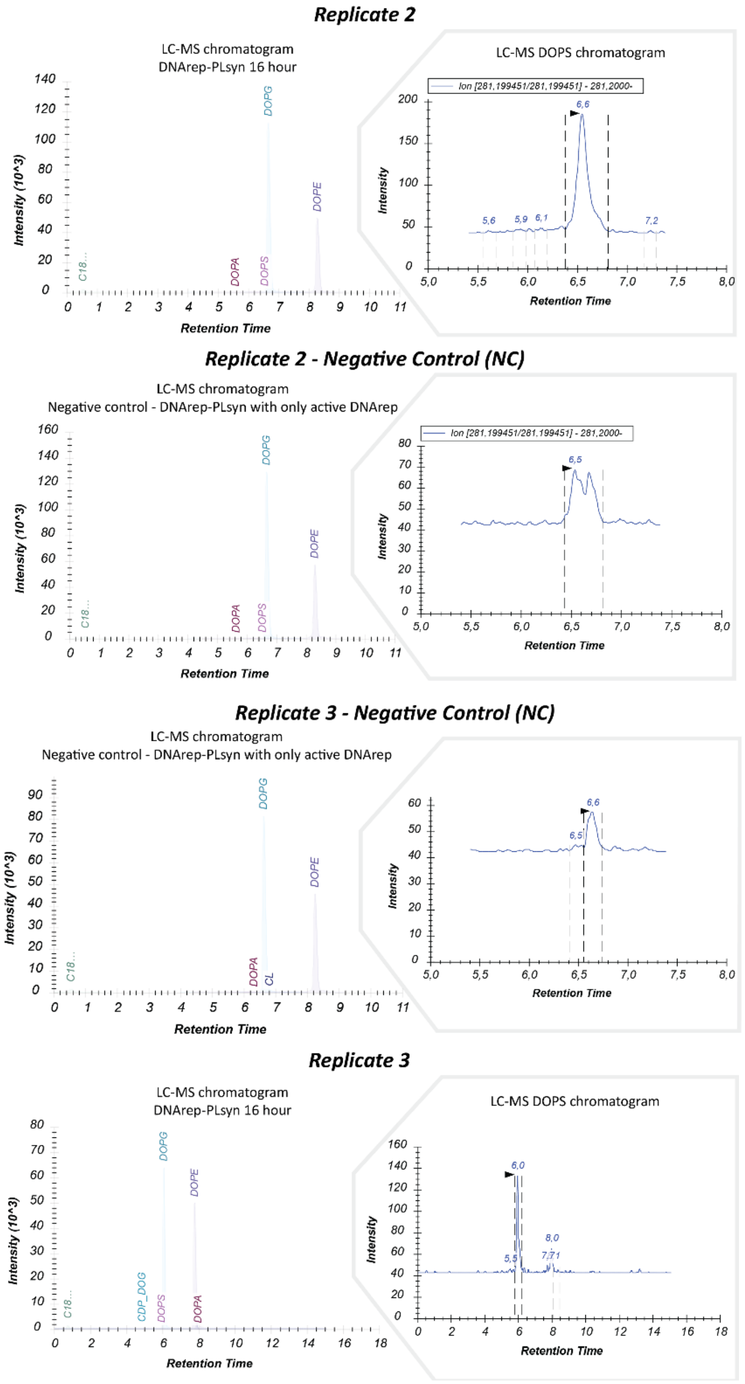


**Fig. S8.** LC-MS chromatograms of the lipid content from liposome samples with expressed *DNArep-PLsyn* and full set of substrates and cofactors. All biological repeats show clear DOPS intensity peaks (insets) when compared to the negative controls (time zero). Intensity values are in arbitrary unit. For biological replicate 1, DOPS peak areas from technical replicates are represented as bar plots.

**Fig. S9.**

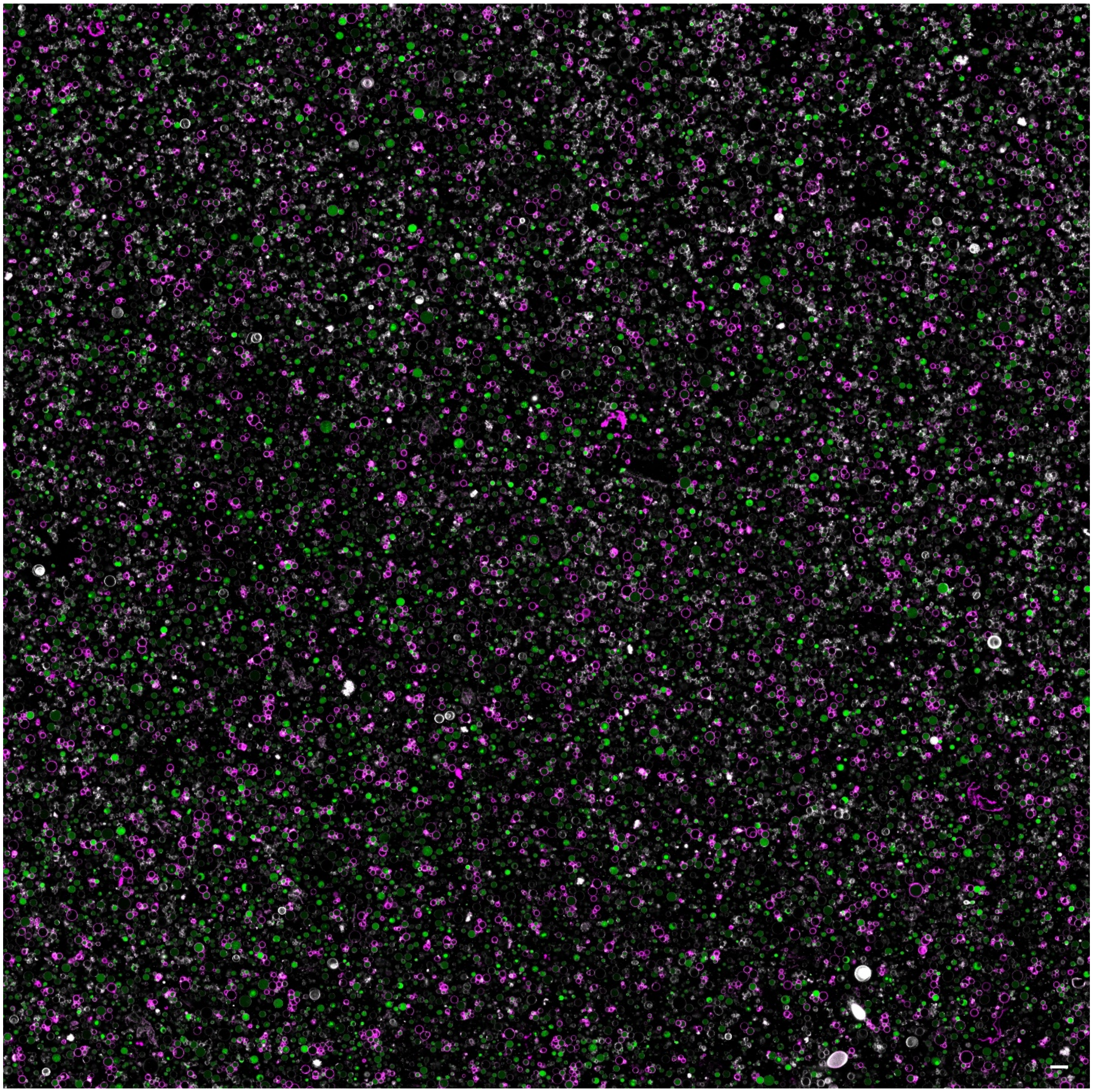

**Fig. S9.** Large field-of-view confocal image (~325x325 µm) from a liposome population expressing *DNArep-PLsyn* with the full set of substrates and cofactors for module activation. White, Cy5 membrane dye; magenta, LactC2-mCherry; green, dsGreen. The scale bar indicates 5 µm. See movie S1 for close-up fields-of-view of the sample.

**Fig. S10.**

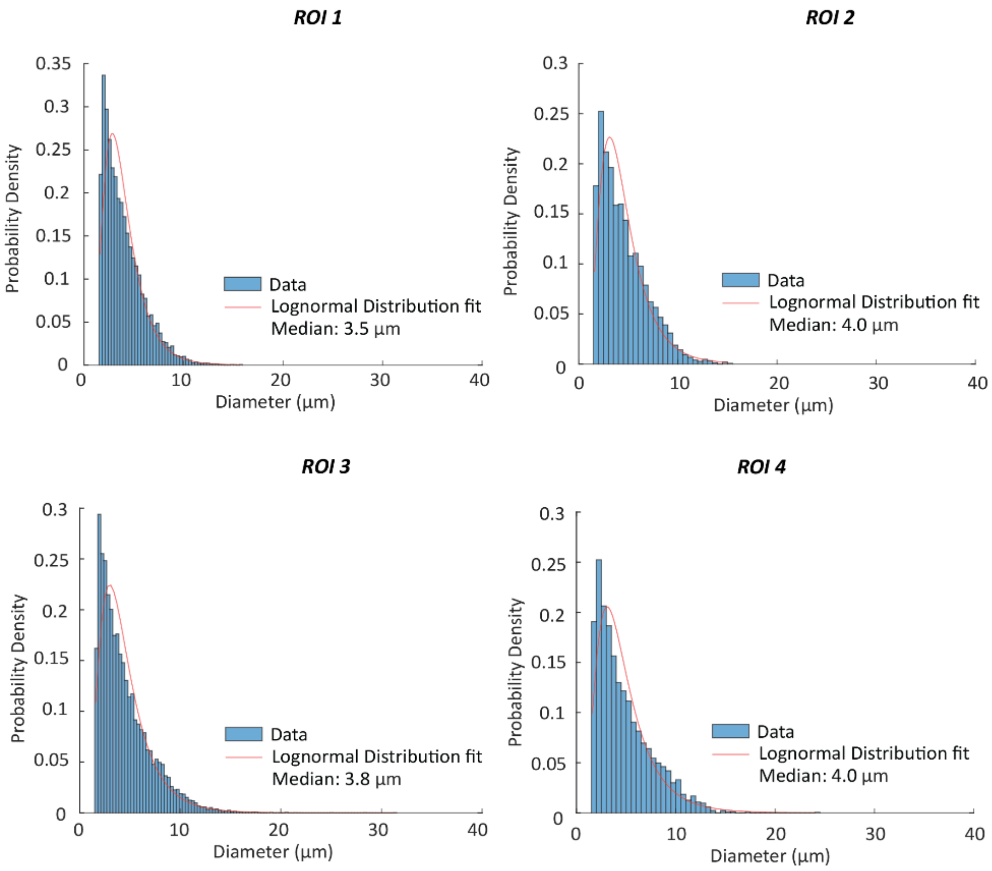


**Fig. S10.** Vesicle size distribution calculated for each ROI from *DNArep-PLsyn*-expressing liposomes. Data from multiple replicate samples were pooled for the analysis. The red curve is the lognormal distribution fit. The median liposome diameter value for each ROI is appended.

**Fig. S11.**

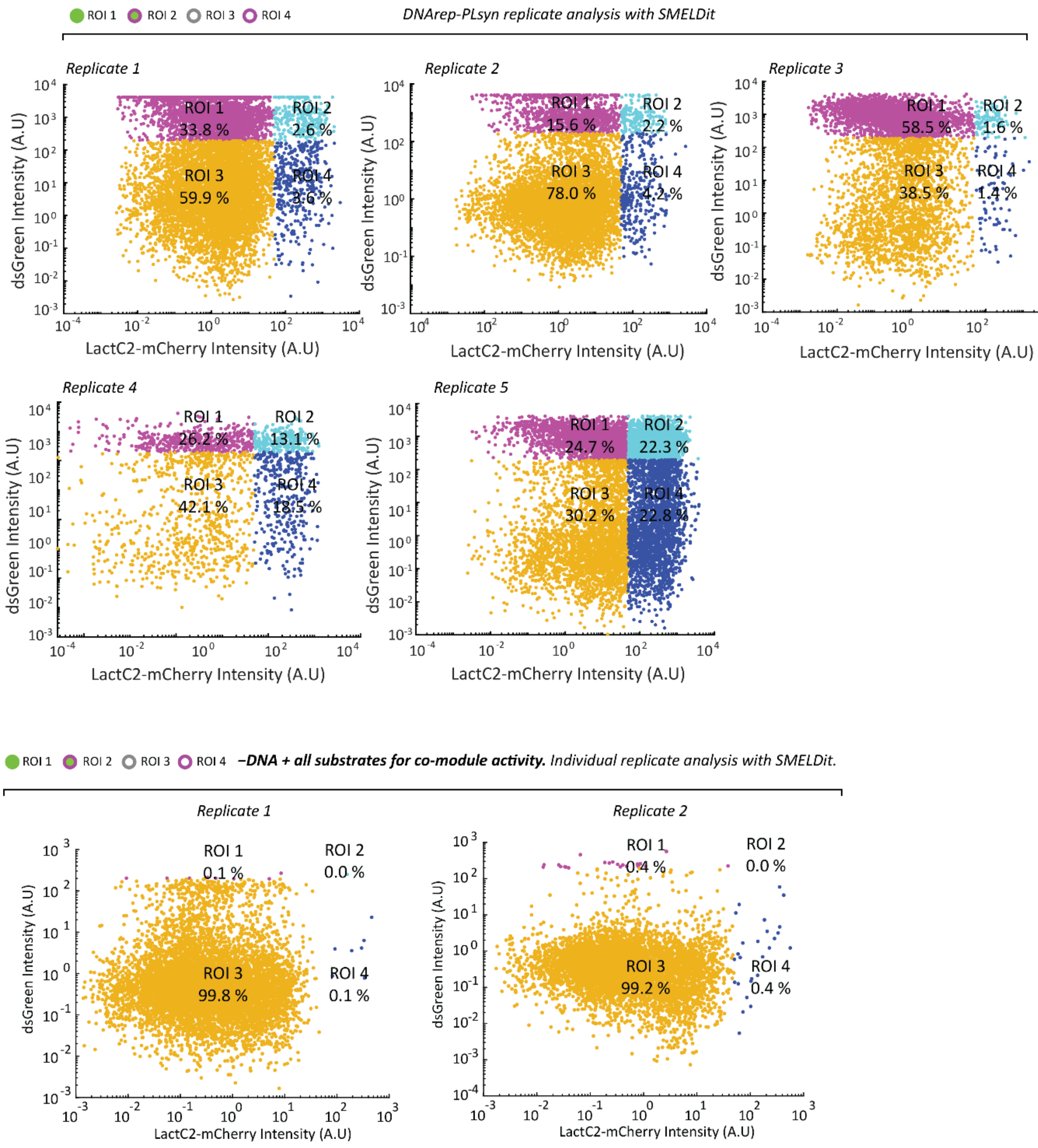


**Fig. S11.** Phenotype scatter plots from SMELDit image analysis (LactC2-mCherry vs. dsGreen) of liposome populations expressing the *DNArep-PLsyn* genome (negative controls with no DNA), in the presence of all substrates and cofactors for dual module activity. Individual biological repeats from pooled data shown in Fig. 3. Displayed ROI percentages were calculated for each replicate. Negative control samples together with experiments shown in Fig. 4 and Fig. 5 were used to define the intensity thresholds for classification into 4 ROIs.

**Fig. S12.**

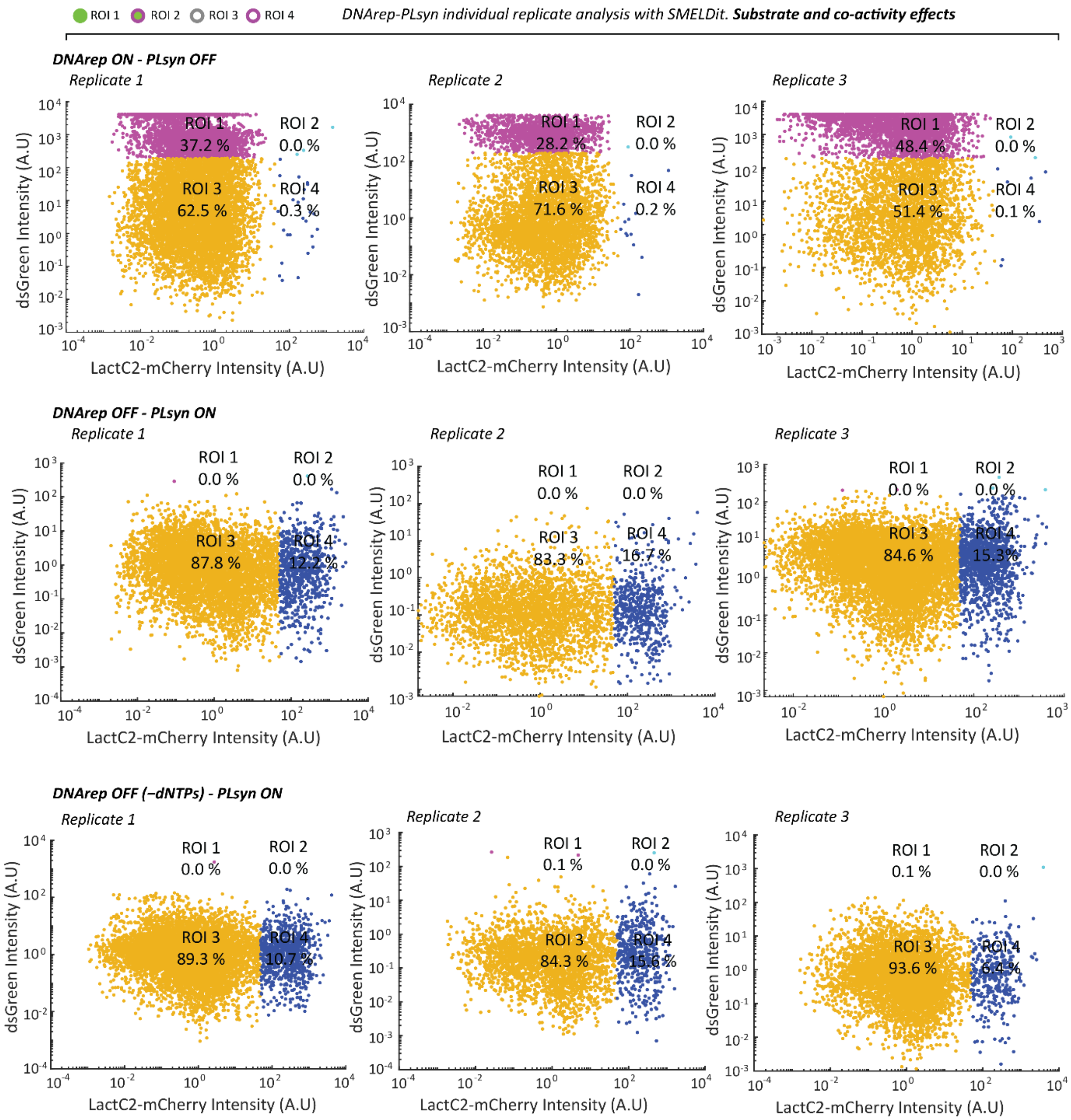


**Fig. S12.** Phenotype scatter plots from SMELDit image analysis (LactC2-mCherry vs. dsGreen) of liposome populations expressing the *DNArep-PLsyn* genome in the presence of substrates and cofactors to activate only DNArep (ON) or only PLsyn (ON). Individual biological repeats from pooled data shown in Fig. 4. Displayed ROI percentages were calculated for each replicate.

**Fig. S13.**

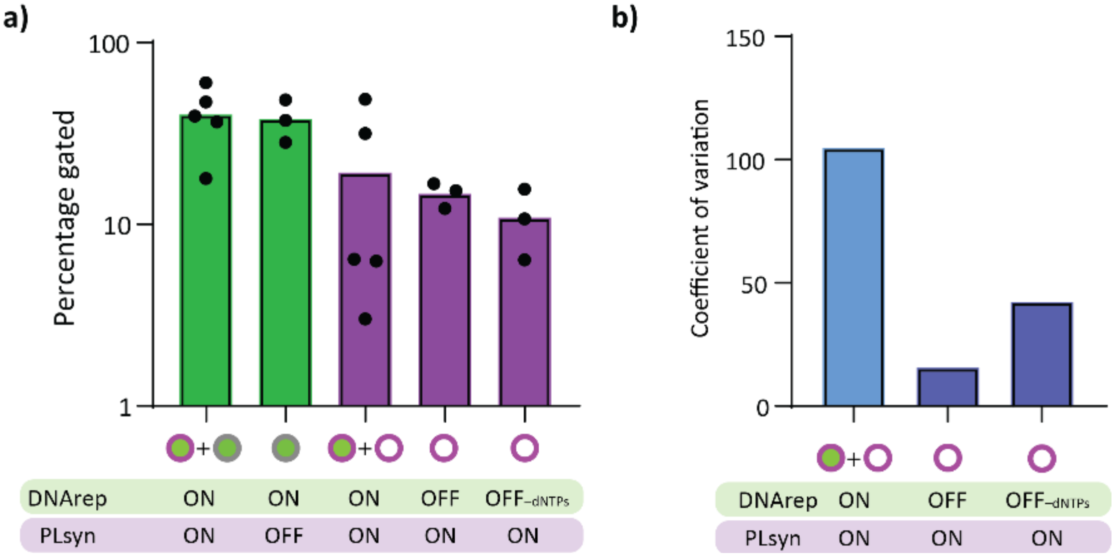


**Fig. S13. a)** Percentage of liposomes expressing the *DNArep-PLsyn* genome with an active DNArep or/and PLsyn module. Data points for each activation condition are from individual biological repeats. **b)** Coefficient of variation calculated from the data shown in a for PLsyn-active conditions.

**Fig. S14.**

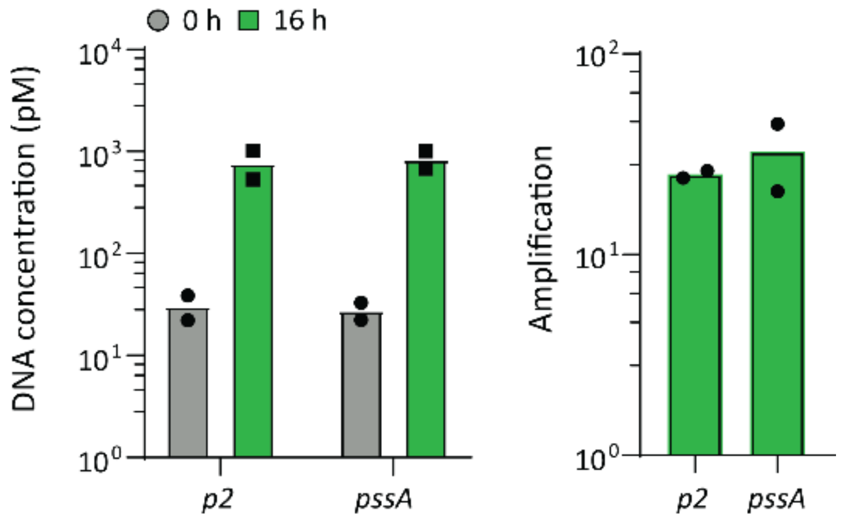


**Fig. S14.** Absolute DNA quantification (left) and replication folds (right) under sole DNArep conditions (PLsyn OFF) resembled those of DNArep-PLsyn reactions with both substrates and cofactors added (main text Fig. 2F). All reactions were performed at 30 °C. The gene names of the targeted regions (~200 bp) are indicated.

**Fig. S15.**

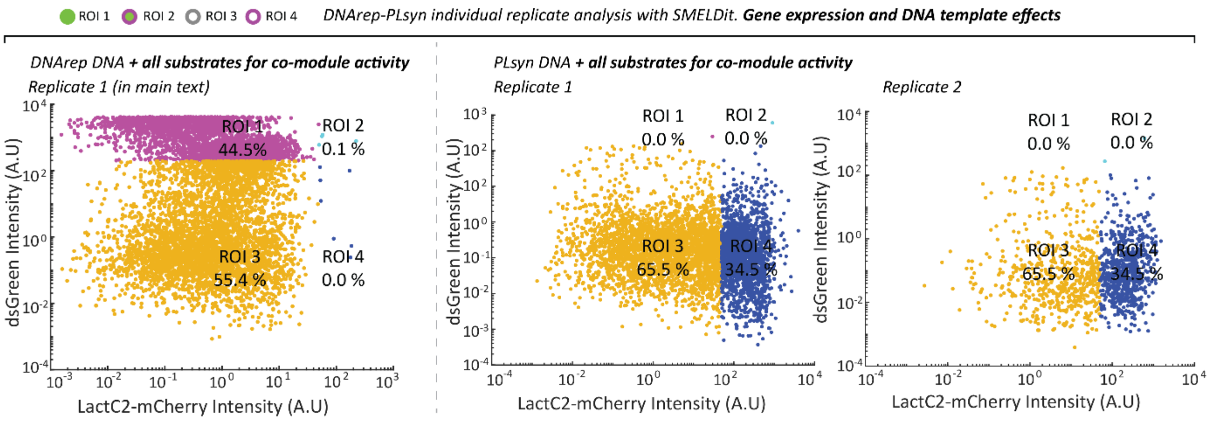


**Fig. S15.** Phenotype scatter plots from SMELDit image analysis (LactC2-mCherry vs. dsGreen) of liposome populations expressing either *DNArep-PLsyn*, *DNArep*, or *PLsyn* DNA in the presence of all substrates and cofactors. Individual biological repeats from pooled data shown in Fig. 5. Displayed ROI percentages were calculated for each replicate.

**Fig. S16.**

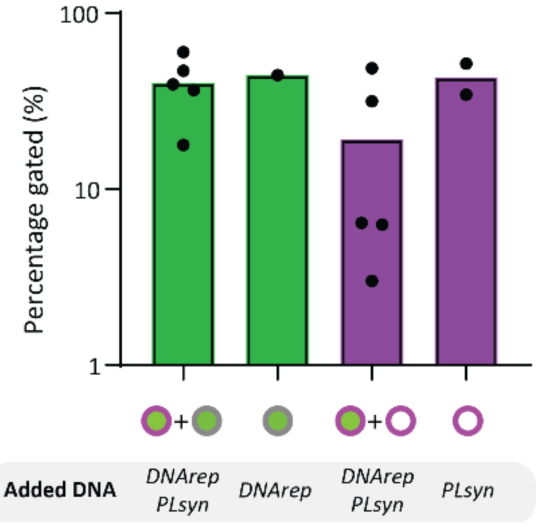


**Fig. S16.** Percentage of liposomes expressing either *DNArep-PLsyn*, *DNArep*, or *PLsyn* DNA in the presence of all substrates and cofactors. Data points for each condition are from individual biological repeats.

**Fig. S17.**

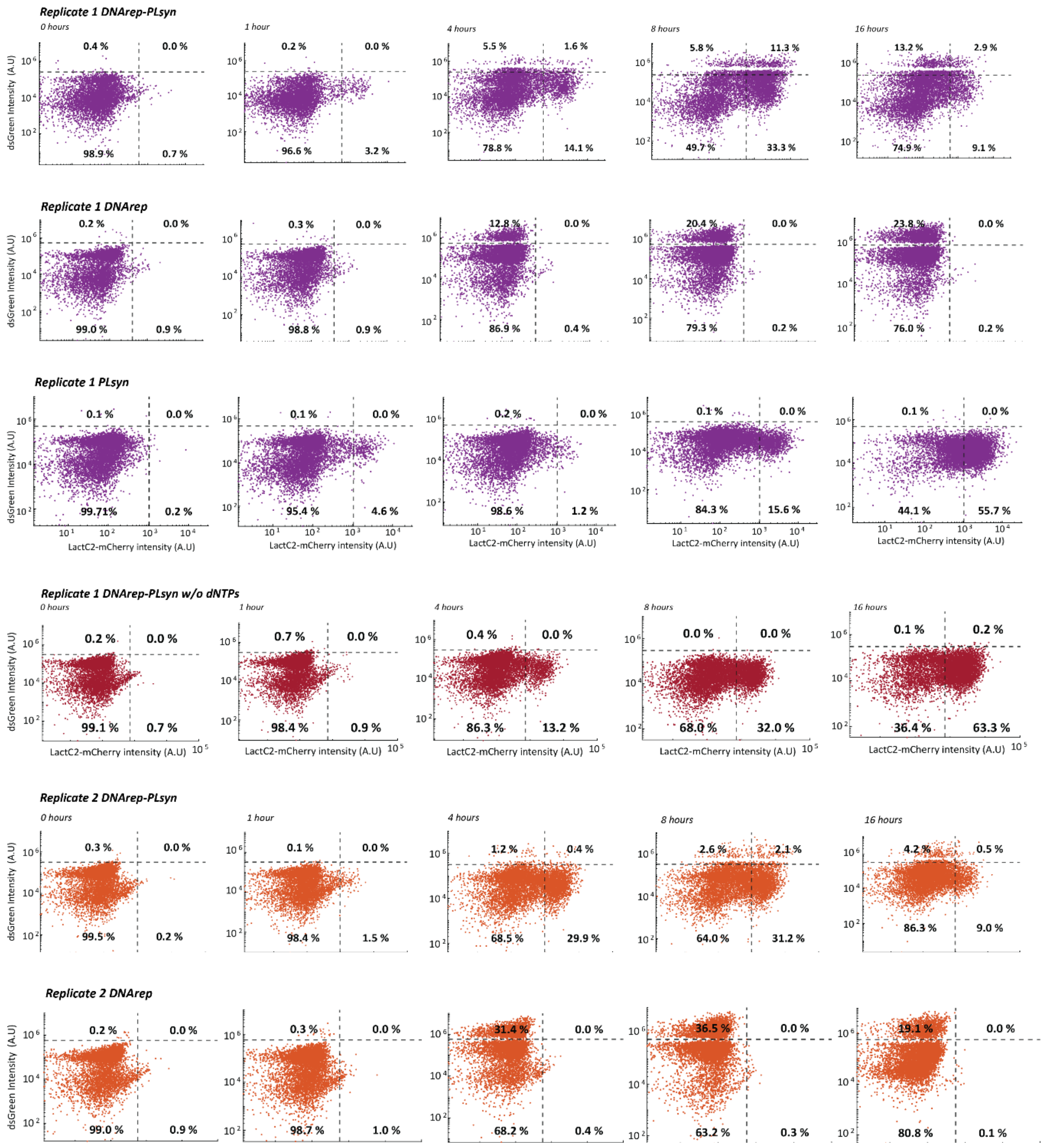


**Fig. S17** Continues next page.

**Fig. S17.**

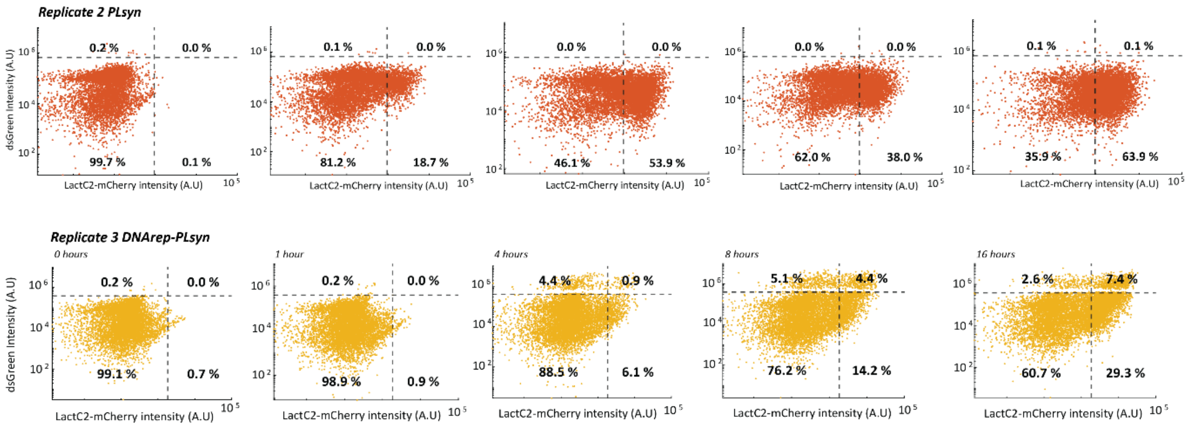


**Fig. S17.** Flow cytometry scatter plots (LactC2-mCherry vs. dsGreen) of liposome samples analyzed at different incubation times for expression of *DNArep-PLsyn*, *DNArep* or *PLsyn* DNA under full substrates/cofactors condition. Data from individual replicate experiments are shown.

**Fig. S18.**

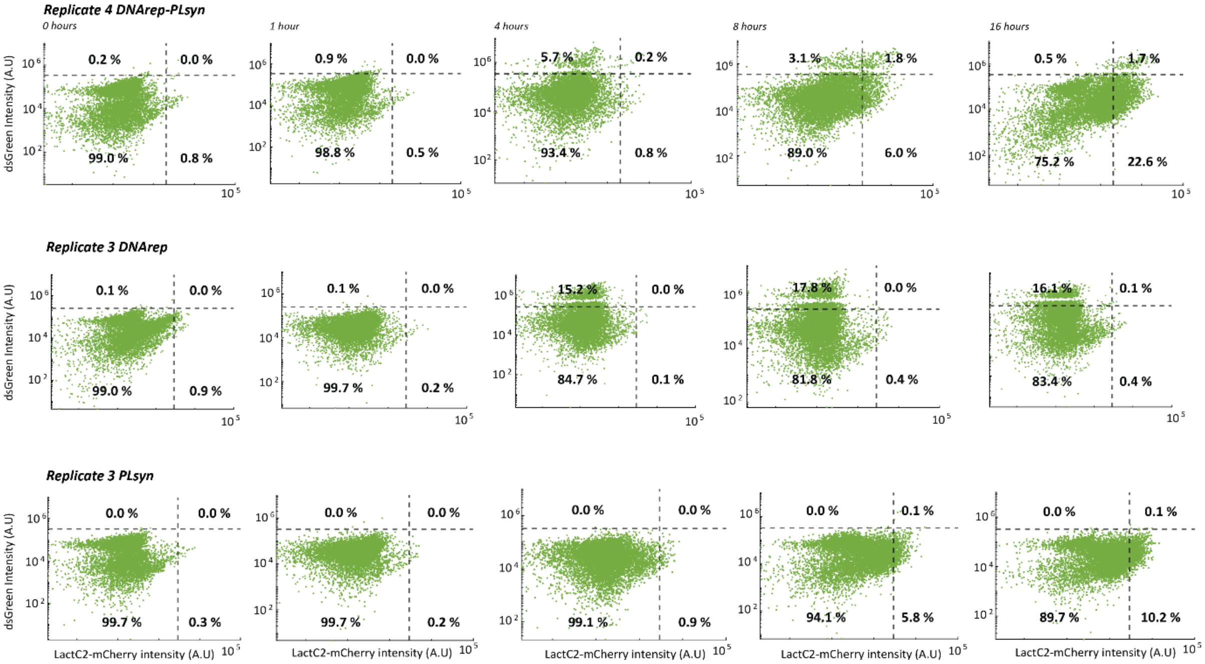


**Fig. S18.** Flow cytometry scatter plots (LactC2-mCherry vs. dsGreen) of liposome samples analyzed at different incubation times for expression of *DNArep-PLsyn*, *DNArep* or *PLsyn* DNA under full substrates/cofactors condition. Contrary to fig. S17, these experiments have been conducted following a slightly different protocol for sample incubation and collection. Here, reactions were incubated in the same tube over all the incubation times. The starting volume of the liposome solution was set to ~10 µL from which three samples of ~2 µL were collected after 1, 4, and 8 hours. The remaining solution was utilized for the 16-hour measurement.

Fig. S19.

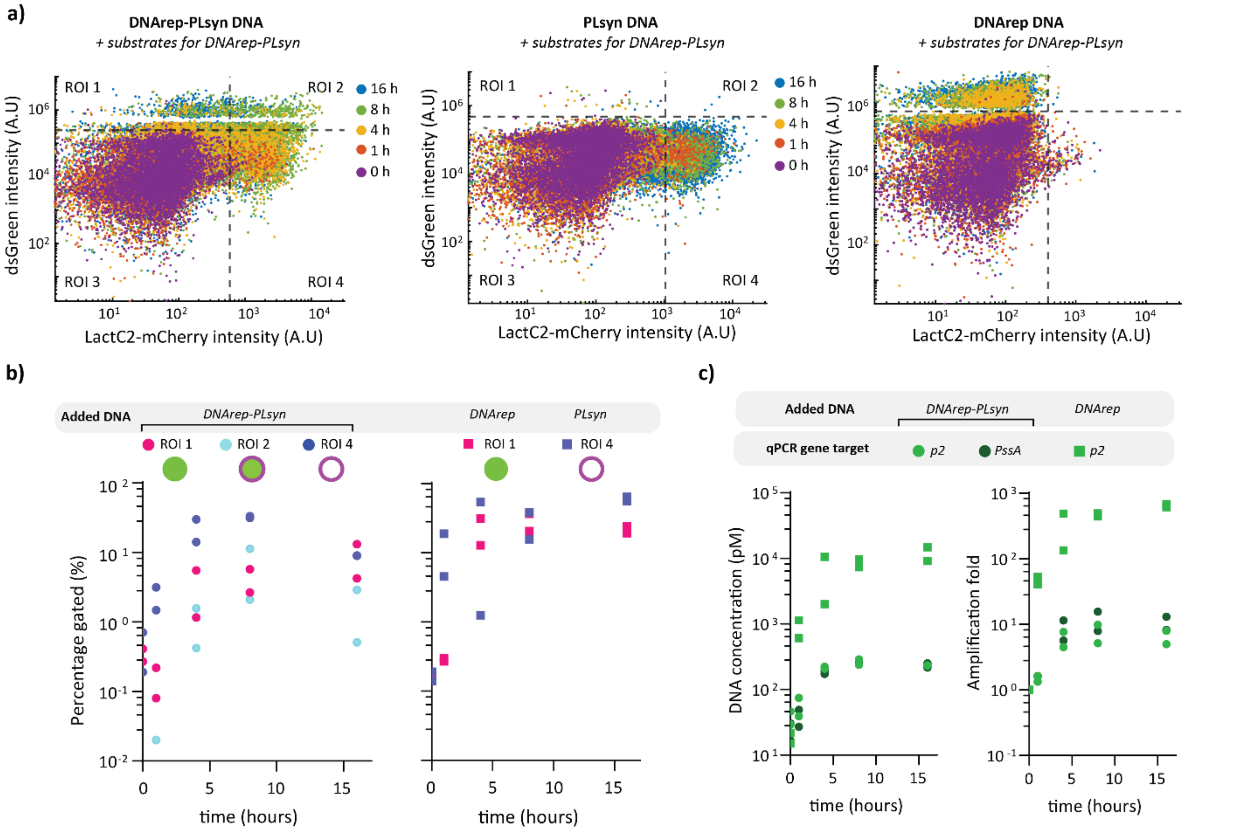


**Fig. S19. In-vesiculo DNArep and PLsyn phenotype appearance along the course of gene expression. a)** Time evolution of flow cytometry scatter plots from liposome samples containing *DNArep-PLsyn*, *DNArep*, or *PLsyn* DNA templates in the presence of all substrates and cofactors. Additional repeats can be found in fig. S17. **b)** Kinetics of the percentage of gated liposomes in the indicated ROIs for the three DNA template conditions. Data points from two biological repeats are displayed. **c)** Kinetics of DNA concentration measured by qPCR from liposome samples containing either *DNArep-PLsyn* or *DNArep* DNA template. Targeted regions for qPCR were located on the *pssA* and *p2* genes of *DNArep-PLsyn*, and on *p2* gene of *DNArep*. DNA amplification fold at different time points was calculated by dividing the concentration of DNA to that at time zero.

**Fig. S20.**

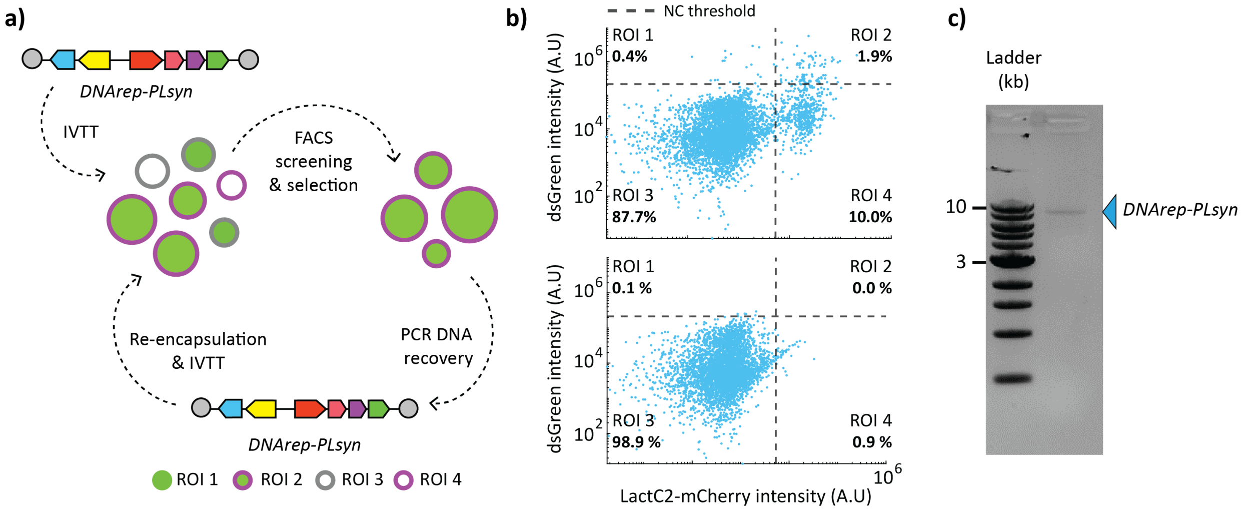


**Fig. S20. Setting up the stage for integrative evolution inside lipid vesicles. a)** Illustration of key experimental steps to perform an evolutionary campaign of *DNArep-PLsyn* inside gene-expressing vesicles: genome (or genome library) encapsulation in liposomes, IVTT, screening and selection through FACS, DNA recovery by PCR, and re-encapsulation to start a next round. **b)** Flow cytometry scatter plots of liposome samples before (0 h, lower panel) and after (16 h, upper panel) expression of 50 pM genome with PURE system. Lipid vesicles exhibiting joint phenotypic traits were gated (ROI 2) and sorted by FACS. **c)** The full-length *DNArep-PLsyn* template can be recovered from the sorted liposomes by PCR, as visualized by agarose gel electrophoresis.

**Table S1.**

| Plasmid name | Plasmid description |
|---|---|
| **G363** | Plasmid encoding for DNArep and PLsyn proteins (DNAP, TP, PlsB, PlsC, CdsA, PssA). All genes are present as individual transcriptional cassettes under the control of an SP6 promoter (DNArep machinery) and T7 promoter (PLsyn machinery), ribosomal binding sites, and T7 transcription terminators. The combined transcriptional units are flanked by Φ29 origin of replication sequences oriL and oriR. |
| **G435** | Plasmid encoding for DNArep proteins (DNAP and TP). Both genes are present as individual transcriptional cassettes, under the control of a T7 promoter, ribosomal binding sites, and T7, vsv-r1 and vsv-r2 transcription terminators (for DNAP and TP, respectively). The combined transcriptional units are flanked by Φ29 origin of replication sequences oriL and oriR. |
| **G555** | Plasmid encoding for PLsyn proteins (PlsB, PlsC, CdsA, PssA). All four genes are present as individual transcriptional cassettes, under the control of a T7 promoter, ribosomal binding sites, and T7 transcription terminators. The combined transcriptional units are flanked by Φ29 origin of replication sequences oriL and oriR. |
| **pRS316** | Plasmid containing the yeast centromeric origin of replication *CEN6/ARS4* and the auxotrophic marker *URA3*. |
| **pY003** | Plasmid assembled in yeast encoding DNArep and PLsyn proteins (DNAP, TP, PlsB, PlsC, CdsA, PssA). All genes are present as individual transcriptional cassettes under the control of a T7 promoter, ribosomal binding sites, and T7 transcription terminators. The combined transcriptional units are flanked by Phi29 origin of replication sequences oriL and oriR. This plasmid also contains the yeast centromeric origin of replication *CEN6/ARS4* and the auxotrophic marker *URA3*. |

**Table S1:** List of plasmids used in this study.

**Table S2.**

| Primer pair | DNA sequence (5' → 3') | Purpose |
|---|---|---|
| **491 ChD**<br>**1302 ChD** | P-AAAGTAAGCCCCCACCCTCACATG<br>GTATTAATTTCACATGCGACAGAATTCGCGGCCGCTTCTAG | PCR amplicon to assemble *DNArep-PLsyn* genome ($PLsyn_{frag}$). DNA template: G363 |
| **1289 ChD**<br>**492 ChD** | GCGAATTCTGTCGCATGTGAAATTAATACGACTCACTATAGGGA<br>P-AAAGTAGGGTACAGCGACAACATACAC | PCR amplicon to assemble *DNArep-PLsyn* ($DNArep_{frag}$). DNA template: G435 |
| **491 ChD**<br>**492 ChD** | P-AAAGTAAGCCCCCACCCTCACATG<br>P-AAAGTAGGGTACAGCGACAACATACAC | PCR to produce *DNArep* and *PLsyn* DNA templates. DNA templates: G435 (for *DNArep*), and G555 (for *PLsyn*). |
| **1459 ChD**<br>**1460 ChD** | CTCCTAATATCGACATAATCCGTCGATCCTCG<br>CCCCATTGACCGACTATCTTCGACAAG | PCR amplicon for one-fragment *DNArep-PLsyn* DNA recovery. 32 bp away from oriR and oriL. |
| **1378 ChD**<br>**1410 ChD** | AAAGTAAGCCCCCACCCTCACATG<br>CGACAGAAACAATCGCACTAAAG | Fragment A for three-fragment *DNArep-PLsyn* DNA recovery. |
| **1411 ChD**<br>**1290 ChD** | CGGGAACATCCAGATGGAAATA<br>ATGCGACAGAATTCGCGGCCGCTTC | Fragment B for three-fragment *DNArep-PLsyn* DNA recovery. |
| **756 ChD**<br>**424 ChD** | ACTTCGCCTTTTTCACGCCC<br>AAAGTAGGGTACAGCGACAACATACAC | Fragment C for three-fragment *DNArep-PLsyn* DNA recovery. |
| **976 ChD**<br>**977 ChD** | GGATGAAGACTACCCGCTGC<br>ACAGGTCTGCGATTTCACCG | qPCR amplicon of targeted region in *p2* gene. |
| **980 ChD**<br>**981 ChD** | ACGGCTGAAATTGACATCCCG<br>CCAGGCGTTGAACTTCTTTGG | qPCR amplicon of targeted region in *p3* gene. |
| **1125 ChD**<br>**1126 ChD** | AACAGGATGACGGTGGCAAA<br>GGAACATCTACGCCCGGATT | qPCR amplicon of targeted region in *pssA* gene. |
| **1119 ChD**<br>**1120 ChD** | TCTCCCGCGACGTATTGATG<br>AATAACGTCCGGCAACTCGT | qPCR amplicon of targeted region in *plsB* gene. |
| **1410 ChD**<br>**1411 ChD** | GGATGAAGACTACCCGCTGC<br>ACAGGTCTGCGATTTCACCG | qPCR amplicon of targeted region in *plsC* gene. |
| **1408 ChD**<br>**1409 ChD** | CGACAGAAACAATCGCACTAAAG<br>CGGGAACATCCAGATGGAAATA | qPCR amplicon of targeted region in *cdsA* gene. |
| **41 ChDT**<br>**58 ChDT** | TTAGGGAGCACATCCATGCCAATAGCTCGACAAGCGGCGAGAG CCTTGCACCTATGCTATATAAAGTAAGCCCCCACCCTCAC<br>GGAAACCGTTGTGGTCTCCCTATAGTGAGTCGTATTAATTTCAC ATGCGACAGAATTCGCGGCCGCTTCTAG | *PLsyn* PCR amplicon to assemble *DNArep-PLsyn* in yeast (pY003). DNA template: G363 |
| **1289 ChD**<br>**44 ChDT** | GCGAATTCTGTCGCATGTGAAATTAATACGACTCACTATAGGGA<br>GCTACATCTTCCGTACTATGCTGTAGTCTCATGGTCGAGTTCTAT TGCTGTTCGGCGGCAAAAGTAGGGTACAGCGACAACATACAC | *DNArep* PCR amplicon to assemble *DNArep-PLsyn* in yeast (pY003). DNA template: G435 |
| **56 ChDT**<br>**57 ChDT** | TGCCGCCGAACAGCAATAGAACTCGACCATGAGACTACAGCAT AGTACGGAAGATGTAGCCCTGGGTCCTTTTCATCACG<br>ATAGCATAGGTGCAAGGCTCTCGCCGCTTGTCGAGCTATTGGCA TGGATGTGCTCCCTAATCTGTGCGGTATTTCACACC | Yeast selection marker (*URA3*) and origin of replication (*CEN6/ARS4*) PCR amplicon to assemble *DNArep-PLsyn* in yeast (pY003). DNA template: pRS316 |
| **50 ChDT**<br>**51 ChDT** | CATGCCAATAGCTCGACAAGC<br>GTACTATGCTGTAGTCTCATGGTCG | Nested PCR (outer PCR) to produce DNArep-PLsyn from yeast isolated DNA. |
| **491E ChDT**<br>**492E ChDT** | P-AAAGTAAGCCCCCACCCTCACATGATACC<br>P-AAAGTAGGGTACAGCGACAACATACACCATTTCC | Nested PCR (inner PCR) to produce DNArep-PLsyn from yeast isolated DNA. |

**Table S2:** List of primers used for PCR and qPCR.

**Movie S1.**

Movie of confocal images from a liposome population expressing *DNArep-PLsyn* with the full set of substrates and cofactors for module activation. The experimental conditions are the same as in Fig. 3 and fig. S9. White, Cy5 membrane dye; magenta, LactC2-mCherry; green, dsGreen.
